# Avirulent ME49Δ*gra5* Protecting Hosts Against *Toxoplasma gondii* Infection and Breast Tumors

**DOI:** 10.1101/2023.02.07.527439

**Authors:** Min Chen, Pei Yang, Zixuan Xin, Jiating Chen, Weihao Zou, Lijuan Zhou, Lili Yang, Jiao Peng, Hongjuan Peng

**Affiliations:** Department of Pathogen Biology, Guangdong Provincial Key Laboratory of Tropical Disease Research, School of Public Health, Southern Medical University, Guangzhou, Guangdong Province, 510515, P. R. China

**Keywords:** *Toxoplasma gondii*, GRA5, vaccine, immune response, 4T1 tumor

## Abstract

*Toxoplasma gondii* is the causative agent of toxoplasmosis, a zoonotic disease that poses a threat to human health and a considerable loss to livestock farming. At present, clinical therapeutic drugs mainly target *T. gondii* tachyzoites and fail to eradicate bradyzoites. Developing a safe and effective vaccine against toxoplasmosis is urgent and important. *T. gondii* possesses dense granule organelles that secrete immunogenic dense granule proteins (GRAs). GRA5 localizes to the parasitophorous vacuole membrane (PVM) in the tachyzoite stage and to the cyst wall in the bradyzoite stage. Here we report that *gra5* knockout *T. gondii* strain is avirulent and fails to form cysts, but stimulates sero-conversion in mice. We next investigated the protective efficacy of ME49Δ*gra5* vaccination against *T. gondii* infection. All the immunized mice survived the challenge infection with either wild-type RH, ME49, VEG tachyzoites or ME49 cysts. Remarkably, ME49Δ*gra5* tachyzoites inoculation *in situ* attenuated the growth of murine breast tumor (4T1) in mice and prevented 4T1’s lung metastasis. ME49Δ*gra5* inoculation might upregulate the levels of Th1 cytokines and tumor-infiltrating T cells in tumor microenvironment (TME) and trigger anti-tumor responses by increasing the number of NK, B, T cells, macrophages, and dendritic cells (DCs) in spleen. Collectively, the results support the potential of ME49Δ*gra5* as attenuated live vaccine against *T. gondii* infection and breast cancer.

**Author summary:** The zoonotic toxoplasmosis poses a great disease burden to humans and big loss to livestock farming. At present, clinical therapeutic drugs mainly target *T. gondii* tachyzoites but have no effect on bradyzoites. A safe and effective vaccine against toxoplasmosis is urgent and important. Dense granule proteins (GRAs) are a group of immunoactive proteins secreted by *T. gondii* dense granule organelles. GRA5 is located on the parasitophorous vacuole membrane (PVM) at the tachyzoite stage and on the cyst wall at the bradyzoite stage. We discovered that the *gra5* knockout *T. gondii* strain became avirulent and failed to form cysts in mice. ME49Δ*gra5* immunized mice survived the challenge infection of wild-type RH, ME49, and VEG tachyzoites, and ME49 cysts. Meanwhile, ME49Δ*gra5* tachyzoites inoculation *in situ* remarkably attenuated the growth of murine breast tumor (4T1) in mice including the injected and distant tumors, as well as prevented 4T1’s lung metastasis. This results support the potential of ME49Δ*gra5* as attenuated live vaccine against *T. gondii* infection and breast cancer.

## Introduction

Toxoplasmosis is a worldwide distributed opportunistic zoonosis caused by the intracellular protozoon *Toxoplasma gondii*. It is estimated that 30% of world human population is infected with *T. gondii* [1].

The parasite secretes dense granule proteins (GRAs) that exhibit high immunogenicity [2]. Upon active host cell entry the fast replicating tachyzoites form a parasitophorous vacuole (PV), in which they reside safely [3]. Bradyzoites are slow growing parasites inside a thick cyst wall and responsible for chronic infection [4]. Protein components of the PV membrane (PVM) and the cyst wall are predominantly released by the dense granule [2, 4]. GRA5 was previously reported to localize to the PVM and the cyst wall [5, 6], however, its function is not known.

Ingestion of raw or under-cooked meat containing *T. gondii* tissue cysts is an important mode of parasite transmission [7]. Although the combination of pyrimethamine and sulfadiazine efficiently kill tachyzoites, the bradyzoites are less impacted due to their very slow growth and the low permeability of the tissue cyst wall in which they reside [8, 9]. Consequently, the development of a safe and effective vaccine is important for the prevention and treatment of toxoplasmosis.

Immunotherapy is very important for cancer treatment and has achieved great success [10]. Among all the immune cells, cytotoxic CD8^+^ T cells have the main effects on prolonging the survival of patients [11]. T helper (T_h_) cells, also known as CD4^+^ T cells, aid the immune activity by releasing cytokines, such as IFN-γ, which are greatly related to the good clinical outcomes for all cancer types [12]. B cells produce antibodies against tumor antigens, which have been frequently found in the serum of cancer patients [13]. Natural killer (NK) cells are a type of cytotoxic lymphocytes critical to the innate and adaptive immune system and kill tumor cells by lysis or inducing apoptosis [11, 14]. Macrophages and dendritic cells (DCs) are innate immune cells in response to tumor antigens [15]. All the above cells are present in the tumor microenvironment (TME). In addition, many processes occur in TME, such as cell apoptosis and immune cell infiltration, are related to cancer immune therapy [16].

Breast cancer (BC) is the most common tumor, as well as the major cause of cancer-related deaths in women all over the world [17]. BC is known as a heterogeneous disease with different pathological features, as BC is a group of distinct neoplastic diseases of the breast tissue and cells, rather than a single disease [18, 19]. Triple-negative breast cancer (TNBC) is one of the BC with the cells deficient for estrogen receptor (ER), progesterone receptors (PR), and human epidermal growth factor receptor 2 (HER2) [20]. Compared to other types of BC, TNBC is characterized by the most aggressive, high recurrence, poor prognosis, and usually accompanied by increased fatality [21]. The treatments for BC include surgery, radiotherapy, and chemotherapy [22]. Among these therapies, chemotherapy is the main therapeutic method for TNBC, however, chemotherapy is prone to induce drug resistance [23]. Because of the high expression of programmed death ligand 1 (PD-L1), TNBC is considered a more immunogenic subtype, therefore immunotherapy has emerged as an important alternative in the management of this cancer [24].

Microorganisms are considered potential cancer immunotherapeutic agents, such as bacteria, viruses, and parasites. The commensal *Bifidobacterium* spp, promote the activity of tumor lymphocytes to suppress B16.SIY melanoma growth [25]. The attenuated *Salmonella* strain (VNP20009) inhibits the growth of primary tumors, and has been used in a phase I study for the patients with metastatic melanoma and renal carcinoma [26]. The attenuated strain of *Listeria monocytogenes* with *actA*/*inlB* deletion improves the survival of mice with ovarian tumor, through iNOS-mediated tumor cell lysis induced by the repolarization of tumor-associated macrophages [27]. The non-pathogenic *E. coli* encoding a nanobody antagonist of CD47 released in the TME through bacteria lysis, activates tumor-infiltrating T cells, stimulates tumor regression, prevents tumor metastasis, and prolongs survival of the mice bearing a syngeneic tumor [28]. Oncolytic viruses target various processes in the cancer-immunity cycle, so they can be used for tumor immunotherapies [29]. Self-assembling virus-like nanoparticles from cowpea mosaic virus (CPMV) are immunogenic, which can inhibit the growth of B16F10 lung melanoma [30]. A combination of CPMV immunotherapy and cyclophosphamide (CPA) inhibited 4T1 murine tumor growth, as well as lung metastasis [31]. *T. gondii* infection induces Th1 immune response that reduces the Lewis lung carcinoma burden [32]. The nonreplicating avirulent uracil auxotroph vaccine strain (*cps*) of *T. gondii* triggers antitumor immune responses against B16F10 melanoma, ovarian cancer, and 4T1 murine cancer [33–35]. Intratumoral injection of the avirulent Δ*gra17* combined with anti-PD-L1 therapy controls growth of murine B16-F10, MC38, and LLC tumors by the synergy effects of these two therapies [36].

In this study, we found GRA5 was critical for *T. gondii* virulence and cysts formation in mice. Vaccination of the ME49 with *gra5* deletion (ME49Δ*gra5)* promoted the levels of IFN-γ, IL12, and TNF-α that protected the mice from challenge infections of wild type RH, ME49 and VEG strains. In addition, ME49Δ*gra5* inoculation not only restrained 4T1 tumors growth but also prevented 4T1’s lung metastasis. Intratumoral injection of ME49Δ*gra5* promoted the production of IFN-γ and IL12 in mice serum and TME, enhanced the anti-tumor responses of spleen, as well as increased the number of tumor-infiltrating T immune cells, such as CD3^+^, CD4^+^, and CD8^+^ cells. Our study supported ME49Δ*gra5* as a promising attenuated vaccine against *T. gondii* infection and as a potential immunotherapy agent for breast cancer.

## Results

### *Tg*GRA5 is critical for *T. gondii* virulence and cyst formation in mice

*Tg*GRA5 previously localizes to the PVM at the tachyzoite stage and lately to the cyst wall at the bradyzoite stage [37]. We further investigated the function of *Tg*GRA5. The *gra5* gene deletion was successfully generated using CRISPR/Cas9 technology (Figure S1A, B). ME49Δ*gra5*’s infection capability was not significantly impaired, compared to which of its parental strain ME49-WT (Figure S1C, D). At 24 h post-infection, the proliferation of ME49Δ*gra5* in HFF cells is modestly inhibited compared to which of ME49-WT. Each PV of ME49-WT contained approximately six tachyzoites on average, while which of ME49Δ*gra5* contained five on average. (Figure 1A, B). In survival assay, the Balb/c mice were intraperitoneal (i.p.) infected with 10^3^ tachyzoites of ME49-WT or ME49Δ*gra5* per mouse (n=6). ME49-WT parasites killed all of the mice at 10 days post-infection (dpi). However, ME49Δ*gra5* parasites exhibited a significant attenuation in virulence, all the mice survived at 30 dpi (Figure 1C). Furtherly, the Balb/c mice were i.p. injected with 10^4^, 10^5^, and 10^6^ ME49Δ*gra5* tachyzoites per mouse (n=6), respectively. We found that the mice challenged with ME49-WT exhibited obvious symptoms such as body shakes, and hair erecting from 6-7 dpi to the death of mice in succession at 10 dpi. However, the mice infected with ME49Δ*gra5* mutant showed very weak symptoms from 6-7 dpi to 14 dpi. The Modified Agglutination Test (MAT) result confirmed the above mice had been successfully infected by ME49-WT or ME49Δ*gra5*, which were all positive with *T. gondii* antibody at 1:12.5 serum dilution (Figure S1E). Only one mouse infected by ME49Δ*gra5* died at the dose of 10^6^ at 7 dpi, and the survival time of the other mice exceeded 30 days (Figure 1C). These results indicated that *Tg*GRA5 played a vital role in the virulence of *T. gondii*. Given that *Tg*GRA5 is a cyst wall protein, we next assessed the ability of ME49Δ*gra5* to establish a chronic infection in mice. Mice were peritoneally injected with 100 tachyzoites of ME49-WT or ME49Δ*gra5*, 75 days later, the mice were sacrificed, and the brains were harvested and homogenized. The brain homogenate smear was examined under a light microscope. We found that ME49-WT parasites formed normal cysts, with 600-800 cysts of about 50 μm in diameter on average in each brain (Figure S1F); remarkably, no cyst was found in the brain homogenate of mice infected with ME49Δ*gra5* (Figure 1D). Moreover, no B1 gene and no BAG1 protein was detected by qPCR or western blot in the brain homogenate of the mice infected with ME49Δ*gra5*, in contrast, both B1 gene and BAG1 protein were detected in brains of the mice infected with ME49-WT at 75 dpi (Figure 1E). To confirm if ME49Δ*gra5* can establish chronic infection or not, the Balb/c mice were further i.p. infected with 10^3^ ME49Δ*gra5* parasites for 7, 15, 30, 75, 150 days, the mice were sacrificed, and the brains were harvested and homogenized, and the B1 gene was detected by q-PCR with the brain homogenate. The B1 gene copy showed the parasite replicated in mice at 15 dpi, but decreased significantly at 30 dpi and to undetectable at 75 and 150 dpi (Figure 1G). The BAG1 protein signal detected by western blot became weaker and weaker from 15 dpi to 30 dpi, and to undetectable at 75 dpi (Figure 1F). These results demonstrated that *Tg*GRA5 is critical for *T. gondii* virulence and cyst formation.

**Figure 1.**
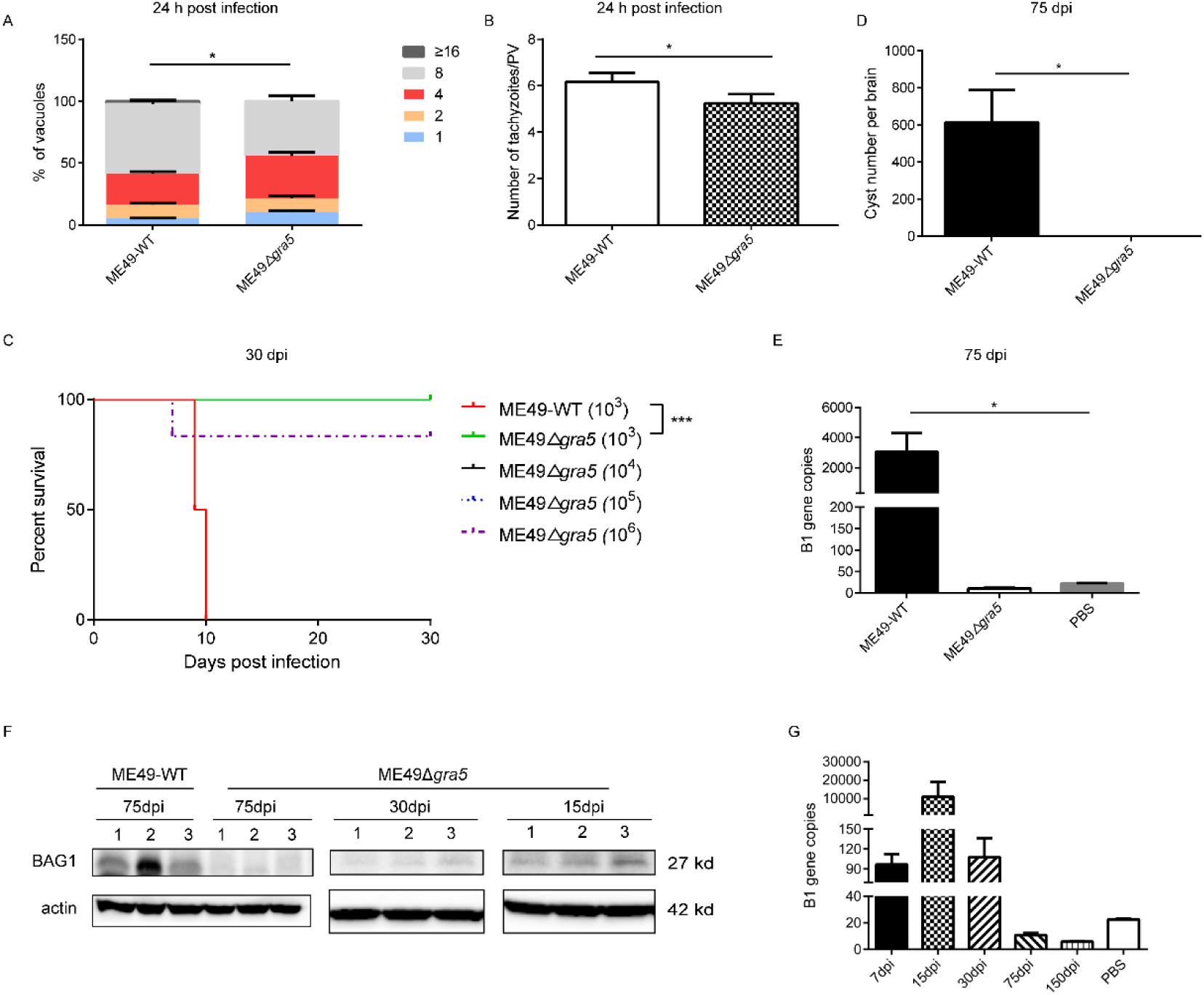
Detection of the virulence and cyst formation of ME49Δ*gra5*. (A-B) HFF cells were infected with ME49-WT or ME49Δ*gra5* tachyzoites at multiplicity of infection (MOI) 1 for 24 h. The number of parasites in each PV (A) and the average number of parasites per PV were calculated (B). (C) 6-8 weeks old Balb/c female mice were i.p. injected with 10^3^ ME49-WT tachyzoites or 10^3^, 10^4^, 10^5^, and 10^6^ ME49-Δ*gra5* tachyzoites respectively, and the survival of mice was monitored for 30 days (n=6 mice). (D-E) 8-week-old male SV129 mice were i.p. infected with 100 ME49-WT or ME49Δ*gra5* tachyzoites (n=3), the mice were sacrificed and each brain was harvested and homogenized at 75 dpi. The cysts of each brain were calculated (D); the B1 copies of brain homogenate was detected by qPCR (E). (F-G) 6-8 weeks old Balb/c female mice were i.p. injected with 10^3^ ME49Δ*gra5* tachyzoites. BAG1 of brain homogenate at 15 dpi and 30 dpi was detected by western blot (F); the B1 copies of brain tissue at 7, 15, 30, 75, 150 dpi was detected by qPCR (G). The Data are represented as mean ± SEM of three independent experiments. Statistical significance was assessed with a two-tailed unpaired Student’s *t*-test (A, B, D, E), log-rank (Mantel– Cox) test (C), or Kruskal-Wallis test (E). \**p*<0.05, \*\*\**p*<0.001.

### The replication of ME49Δ*gra5* strain is inhibited in mice

We further investigated whether *Tg*GRA5 is essential for the proliferation of *T. gondii in vivo*. Balb/c Mice were i.p. injected with 10^3^ tachyzoites of ME49-WT or ME49Δ*gra5*. The weight of mice was recorded before infection and at 7 dpi. The result showed that the weight gain of the mice infected with ME49-WT was significantly less than that of the mice infected with ME49Δ*gra5* and the PBS control group; no significant difference in the body weight changes was found between the ME49Δ*gra5* infection group and the PBS control group (Figure S2A). In addition, the mice were sacrificed and dissected at 7 dpi, and we found that the weight of the spleen and liver of the mice infected with ME49-WT and ME49Δ*gra5* was significantly higher than that of the PBS control group, but no significant difference in the weight of brain and lung was found among these three groups (Figure S2B-E). We next explored whether ME49Δ*gra5* strain has replication defect in mice by detecting the parasitic load in different organs. At 7 dpi, the mice were sacrificed, and the peritoneal fluid, spleen, liver, brain, and lung were harvested and subjected to qPCR to detect B1 gene copies. The parasitic load in peritoneal fluid, spleen, liver, brain, and lung in the mice infected with ME49Δ*gra5,* was significantly lower than that of the mice infected with ME49-WT (Figure S3A-E). These results suggest that the proliferation of *T. gondii in vivo* is significantly inhibited when *gra5* gene is deleted.

### ME49Δ*gra5* stimulates immune responses in mice

To assess the role of the immune responses which led to survival of mice infected by ME49Δ*gra5*, Balb/c mice were i.p. injected with 10^3^ tachyzoites of ME49-WT, ME49Δ*gra5* or PBS as the control. At 7 dpi, the spleen of mice was collected for RNA sequencing. Differentially expressed genes (DEGs) were selected according to | log_2_FC |≧1 (FC, fold change) and *p*-adjust<0.05. Compared with the PBS treatment group, the up-regulated genes of ME49-WT or ME49Δ*gra5* infected mice were subjected to KEGG and REACTOME analysis. The results showed that in ME49-WT group, the most up-regulated genes were both enriched in the immune responses (Figure S4A, B) as well as in the mice infected with ME49Δ*gra5* (Figure S4C, D). We further analyzed the DEGs between the ME49-WT and ME49Δ*gra5* infected mice, and found 148 DEGs, of which, 56 genes were up-regulated and 92 genes were down-regulated (Figure 2A). The heat map of DEGs showed that most of these genes were related to immunity and signal transduction pathways (Figure 2B). To further dissect these DEGs, the top 24 DEGs were subjected to GO and KEGG enrichment analysis. GO enrichment showed that the 24 DEGs were related to inflammatory response (Figure 2C). KEGG enrichment showed that the pathways enriched by most of the DEGs were related to immunity, including cytokine and cytokine receptor interaction, chemokine signaling pathway including IL-17 signaling pathway and TNF signaling pathway. (Figure 2D).

**Figure 2.**
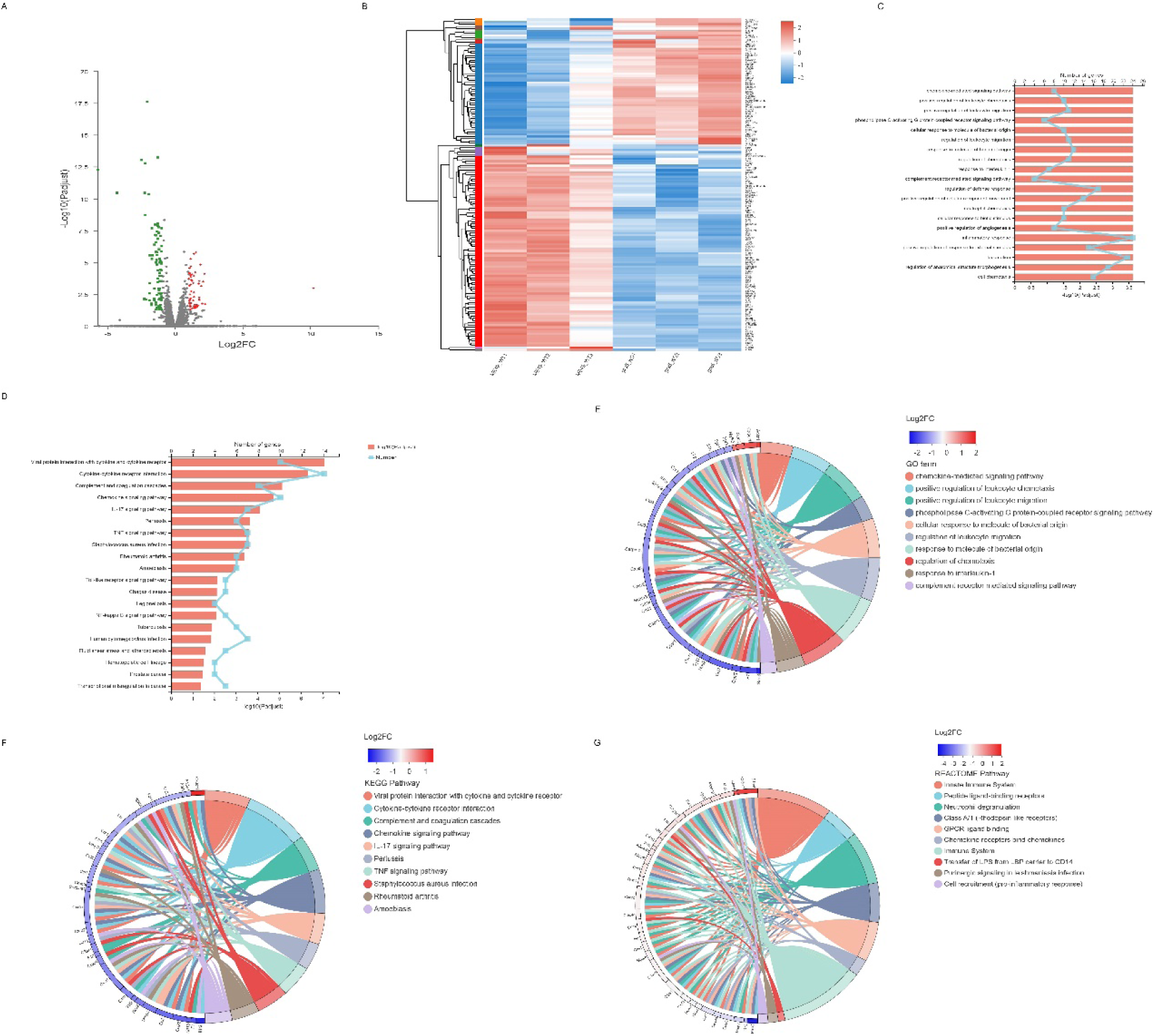
Bioinformatic Analysis for differentially expressed genes (DEGs) of murine spleen infected with ME49-WT or ME49Δ*gra5* parasites. 6-8 weeks old female Balb/c mice were injected with 10^3^ ME49-WT, ME49Δ*gra5* parasites or PBS for 7 days. The spleen RNA was collected for RNA sequencing. (A) 56 DEGs were up-regulated and 92 DEGs were down-regulated for ME49-Δ*gra5* infected mice, compared with ME49-WT. (B) The heat map of DEGs. (C) GO enrichment of DEGs. (D) KEGG enrichment of DEGs. (E-D) Enrichment chord analysis of DEGs. GO term (E), KEGG pathway (F), REACTOME pathway (G).

Furtherly, the GO enrichment chord diagram showed that most of the DEGs were down-regulated, in which the most down-regulated gene, Saa3, was enriched in response to interleukin-1 (Figure 2E). KEGG pathway enrichment chord diagram indicated that Cx3cr1 expression was significantly increased, which was mainly involved in the interaction of viral proteins with cytokines and cytokine receptors, cytokine-cytokine receptor interaction, and chemokine signaling pathway. In addition, the significantly reduced expression of IL1r2 was mainly related to the cytokine-cytokine receptor interaction and the Amoebiasis (Figure 2F). The results of the REACTOME enrichment chord showed that there were 25 DEGs involved in adaptive immunity pathways, and 22 DEGs involved in innate immunity (Figure 2G).

### ME49Δ*gra5* vaccination protects mice from tachyzoites and bradyzoites infection

Due to its low virulence, defect in cysts formation, and stimulation of immune responses in mice, ME49Δ*gra5* was assessed as a vaccine candidate. Balb/c mice were immunized with 10^3^ ME49Δ*gra5* tachyzoites i.p. At 30 dpi, both the vaccinated and the naïve mice were challenged with 10^3^ tachyzoites of WT RH, ME49, or VEG, and were monitored for another 30 days. All non-immunized mice died within 11 days after infection. while all ME49Δ*gra5* vaccinated mice survived to the end of the experiment after infection by these three types of *T. gondii* (Figure 3A-C). We further detected whether ME49Δ*gra5* vaccination produced long-term immune protection against *T. gondii* challenging infection. Balb/c mice were immunized with 10^3^ ME49Δ*gra5* tachyzoites for 75 days or unimmunized, and then they were injected i.p. with 10^3^ tachyzoites of RH-WT, ME49-WT, or VEG-WT for another 30 days. The survival rates of the immunized mice were 100%, while the naïve mice all died within 9 days after infection (Figure 3D-E).

**Figure 3.**
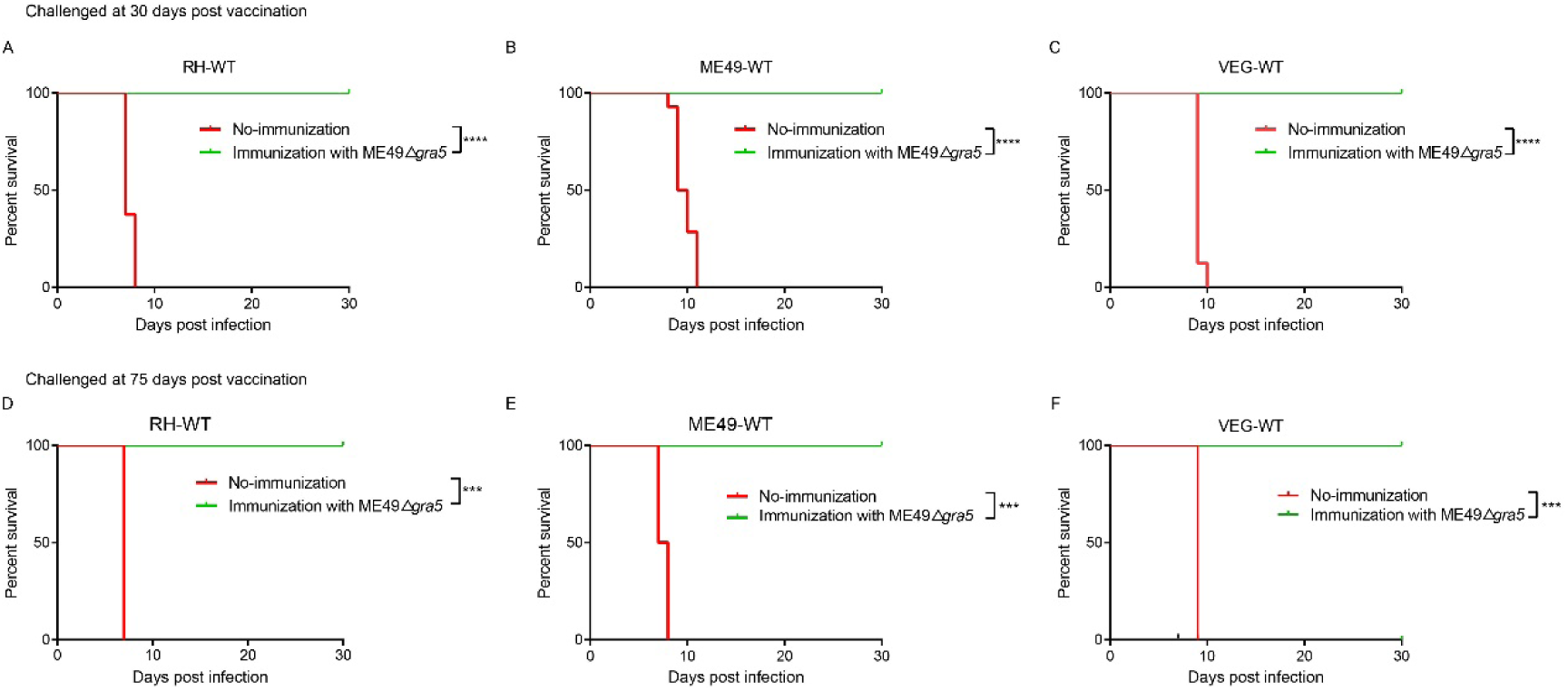
Examination of the protective effects of ME49Δ*gra5* vaccination against different types of *T. gondii*. (A-F) The survival curves of Balb/c mice after infection by different types of *T. gondii* strains as indicated. 6-8 weeks old female Balb/c mice were immunized with 10^3^ ME49Δ*gra5* parasites for 30 days (A, B, C) or 75 days (D, E, F), and then the mice were challenged with 10^3^ tachyzoites of RH-WT (A, D), ME49-WT (B, E) or VEG-WT (C, F). All mice were monitored for 30 days, and the non-vaccinated mice were used as control (n=8). Data are represented as mean ± SEM of three independent experiments. Statistical significance was assessed with log-rank (Mantel–Cox) test. \*\*\**p*< 0.001, \*\*\*\**p*<0.0001.

Since ME49Δ*gra5* vaccination had a significant protective effect on *T. gondii* tachyzoite infection, we tested whether it also could effectively resist bradyzoites infection. Thirty days after the immunization with 10^3^ ME49Δ*gra5* tachyzoites, the mice were orally challenged with 20 cysts of ME49-WT. The result showed that all vaccinated mice survived the cyst challenge to 30 dpi. However, non-immunized mice all died within 10 days after infection (Figure S5). These results demonstrate that ME49Δ*gra5* strain can provide strong immune protection against tachyzoites and bradyzoites infection.

### ME49Δ*gra5* vaccination produces pro-inflammatory cytokines

To evaluate the potential mechanism of immune protection of ME49Δ*gra5* strain, the levels of pro-inflammatory cytokines were measured in mice serum by ELISA at 30 days or 75 days after vaccination. IFN-γ, IL12, and TNF-α are the key cytokines against *T. gondii* infection. Therefore, we detected the levels of these cytokines. The results showed that IFN-γ, IL12, and TNF-α level was significantly elevated at both 30 and 75 days post-immunization, compared to that of the naïve mice (Figure 4A-C).

**Figure 4.**
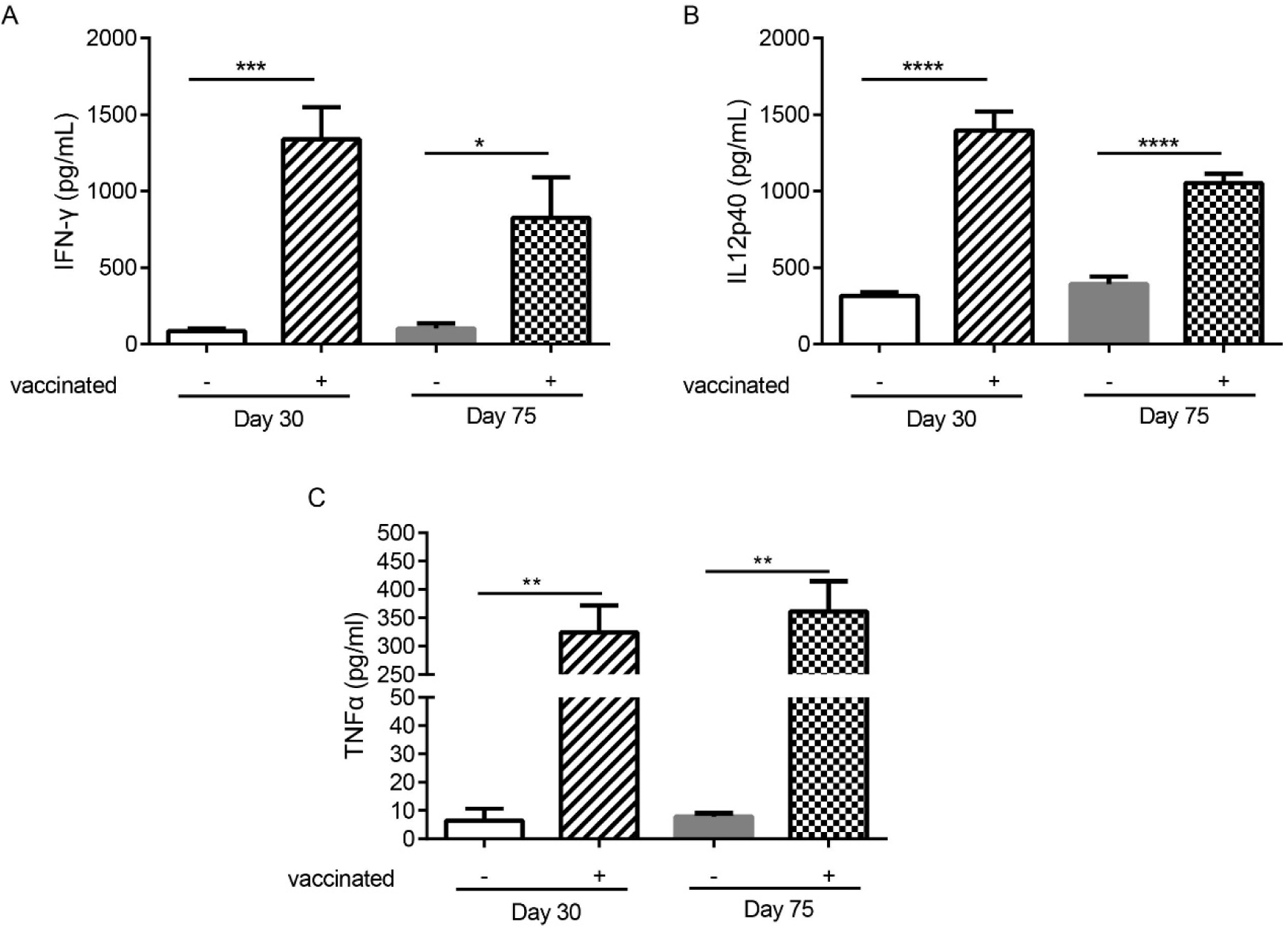
ME49Δ*gra5* parasites vaccination inducing host immune responses. (A-C) 6-8 weeks old female Balb/c mice were immunized with 10^3^ ME49Δ*gra5* parasites for 30 or 75 days. The serum of immunized and naïve mice (n=5) was obtained and detected by ELISA for the levels of IFN-γ (A), IL12p40 (B), and TNF-α (C). Data are represented as mean ± SEM of three independent experiments. Statistical significance was assessed with a two-tailed unpaired Student’s *t*-test. \**p*<0.05, \*\**p*<0.01, \*\*\**p*<0.001, \*\*\*\**p*<0.0001.

### ME49Δ*gra5* suppresses 4T1 tumor growth

It has been previously reported that the attenuated *T. gondii* inhibits tumor growth and metastasis [35, 36, 38]. As the avirulent ME49Δ*gra5* tachyzoites could prominently induce pro-inflammatory cytokines, we then continued to determine its antitumor effects with 4T1 murine breast tumor as the model. The tumor was inoculated *in situ* with ME49Δ*gra5* tachyzoites on day 9, 11, and 13 post 4T1 cells injection, with a dose of 10^5^ tachyzoites per inoculation (Figure 5A). The mice with intratumoral inoculation of ME49Δ*gra5* showed remarkable tumor growth inhibition, including the significantly decreased weight and size of the tumors, compared with that of the PBS inoculation (control) group (Figure 5B, C, D). On day 27 post tumor inoculation, the tumor volumes of the control group were six-fold larger than that of the ME49Δ*gra5* inoculation group (Figure 5D). ME49Δ*gra5* inoculation significantly extended the survival of the mice bearing 4T1 tumor, compared with the PBS treatment (Figure 5E).

**Figure 5.**
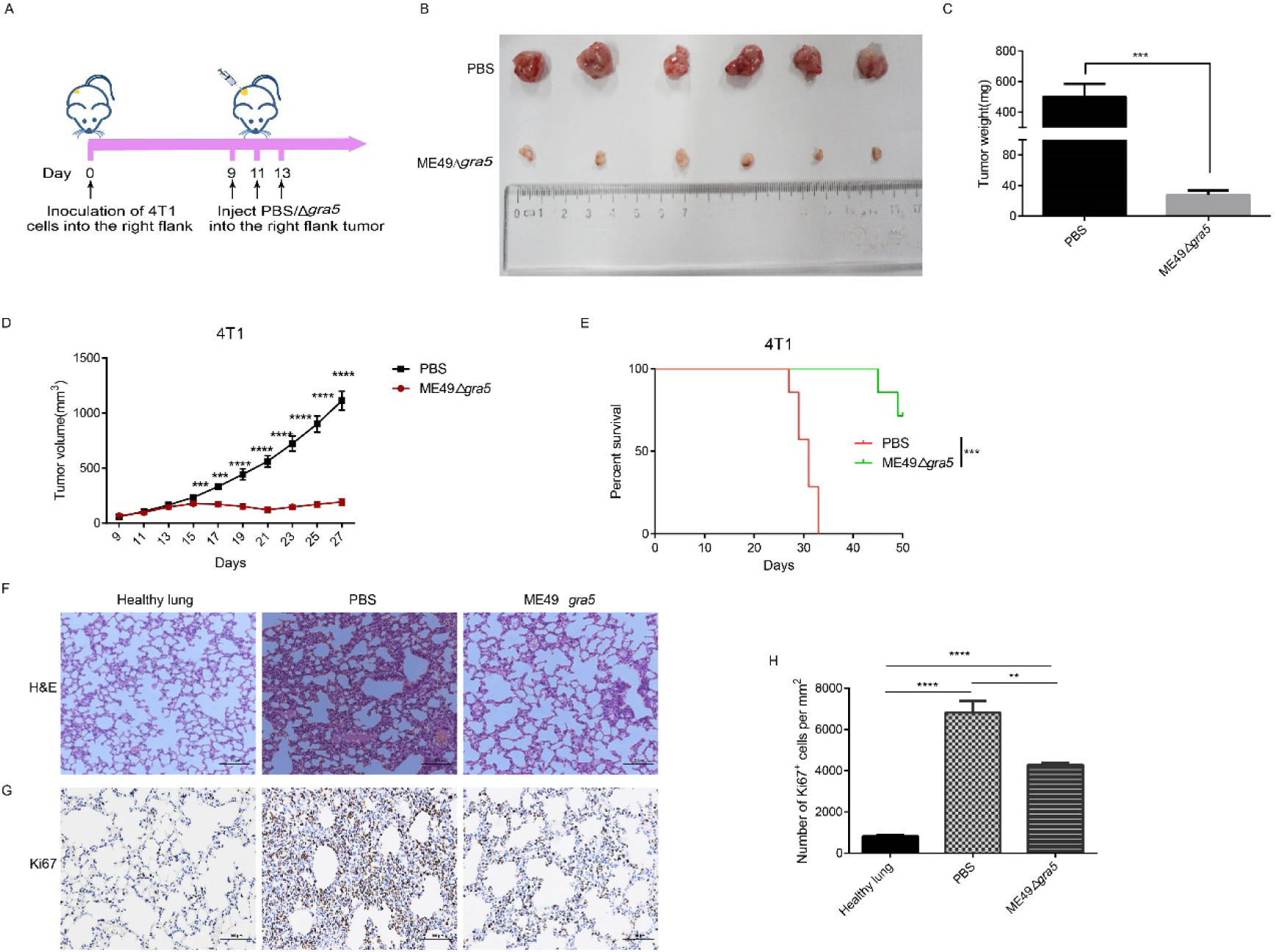
ME49Δ*gra5* tachyzoites treatment inhibited 4T1 tumor growth in mice. (A) Schematic diagram to show the processes of mice being inoculated with 4T1 cells and treated with ME49Δ*gra5* tachyzoites or PBS. (B-H) Comparisons between the 4T1 tumor bearing mice treated with ME49-Δ*gra5* or PBS: the tumors (B); tumor size (C); tumor volume (D); survival curves (E); lung sections stained by H&E staining (F); immunohistochemistry analysis of lung tissue stained with Ki67 antibody (G); The abundance of Ki67-positive cells in the lung tissue (H). Data are represented as mean ± SEM of three independent experiments (A-E, n=7, F-H, n=3). Statistical significance was assessed with a two-tailed unpaired Student’s *t*-test (D), log-rank (Mantel–Cox) test (E), or One-way ANOVA (H). \*\**p*<0.01, \*\*\**p*<0.001, \*\*\*\**p*<0.0001. Scale bar=100 μm.

4T1 primary tumor can develop spontaneous metastatic disease in the lungs of mice as early as 8 days after inoculation [31, 35]. We continued to evaluate if ME49Δ*gra5* inoculation can inhibit 4T1 metastasis. The lung sessions with H&E staining showed that compared with the PBS group, the infiltrating myeloid cells were significantly reduced, and the alveolar spaces were larger in the ME49Δ*gra5* inoculation group (Figure 5F). We also assessed 4T1 lung metastasis by using Ki67 antibody to stain the lung tissue. Less Ki67-positive cells was detected in the lungs of the mice inoculated with ME49Δ*gra5,* compared with that in the control group (Figure 5 G, H). Based on these results, we conclude that ME49Δ*gra5* parasites treatment inhibits 4T1 tumor growth and lung metastasis.

### ME49Δ*gra5* treatment increases tumor-infiltrating lymphocytes (TILs)

Cytokines secreted by immune cells play crucial roles in suppressing tumor progression. IL12 is the primary driver of Th1 differentiation and stimulates the production of IFN-γ to coordinately attack tumors [31]. Therefore, we performed ELISA assay to estimate the levels of IFN-γ and IL12 in the serum and homogenized tumor lysate of the mice after ME49Δ*gra5* inoculation. We found that ME49Δ*gra5 in situ* inoculation increased the level of IFN-γ and IL12 in both mice serum and the tumor microenvironment (TME), compared to PBS inoculation (Figure S6). Cytokines activate immune cells and induce their concentration to kill tumor cells. As expected, ME49Δ*gra5* therapy increased the population of T cells, as indicated by the increased CD3+ (Figure 6A, B), CD4+ (Figure 6C, D), and CD8+ cells (Figure 6E, F). Taken together, ME49Δ*gra5* treatment elevates the levels of antitumor cytokines and the amounts of tumor-infiltrating lymphocytes.

**Figure 6.**
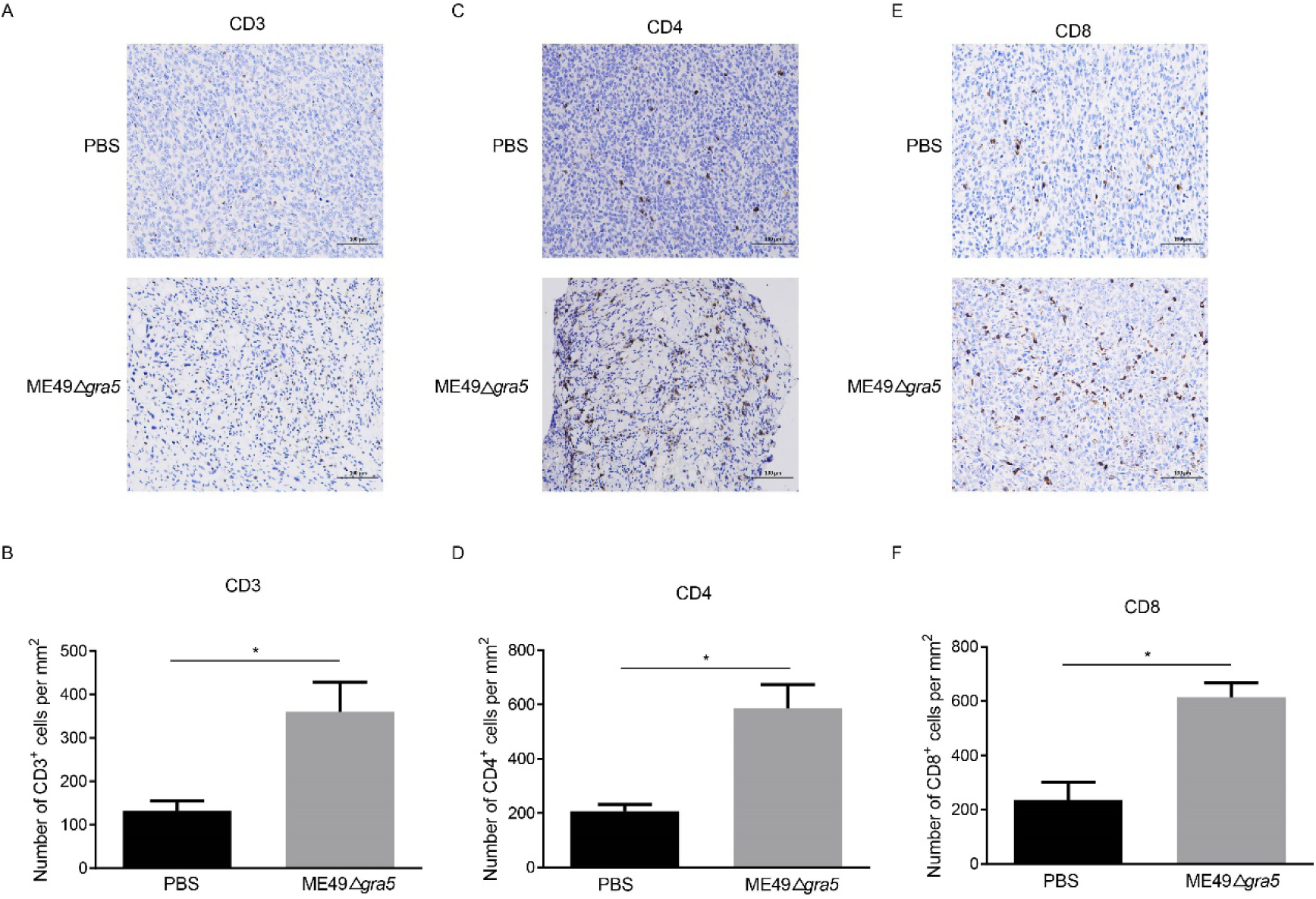
ME49-Δ*gra5* tachyzoites inoculation increased tumor-infiltrating lymphocytes (TILs) in mice. (A-F) 4T1 tumors were harvested on day 21 post 4T1 cell inoculation and used for immunohistochemistry staining for CD3 (A), CD4 (C), and CD8 (E). The numbers of CD3+ (B), CD4+ (D), and CD8+ (F) cells were calculated and compared between the PBS and the ME49-Δ*gra5* inoculation groups. Data are represented as mean ± SEM of three independent experiments (n=3). Statistical significance was assessed with a two-tailed unpaired Student’s *t*-test. \**p*<0.05. Scale bar=100 μm.

### ME49Δ*gra5* treatment enhances the anti-tumor immune responses of spleen

We further analyzed the immune status of spleen at 21 dpi. The innate immune cells, CD49b, F4/80, and CD11c representing NK cells, macrophages, and DCs respectively, were remarkably increased in ME49Δ*gra5* treated mice (Figure 7A, B, F, G). CD8^+^ T cells are the major cell type mediating anti-tumor effects, and the CD8^+^/CD4^+^ ratio indicates the amount of CD8^+^ T cells of specific anti-tumor immunity [39]. We found that the amount of CD8^+^ T cells and the ratio of CD8^+^/CD4^+^ were significantly elevated in ME49Δ*gra5* treated mice (Figure 7A, C, D). The production of CD4^+^ T follicular helper cells requires B cells that recognize tumor neoantigens and their collaboration promotes anti-tumor CD8^+^ T cells responses [40]. Concordantly, the quantity of B cells marked by CD19 (a marker of B cells), was significantly increased in ME49Δ*gra5* therapy group (Figure 7A, E). The above results were consistent with the immune responses in TME (Figure 6).

**Figure 7.**
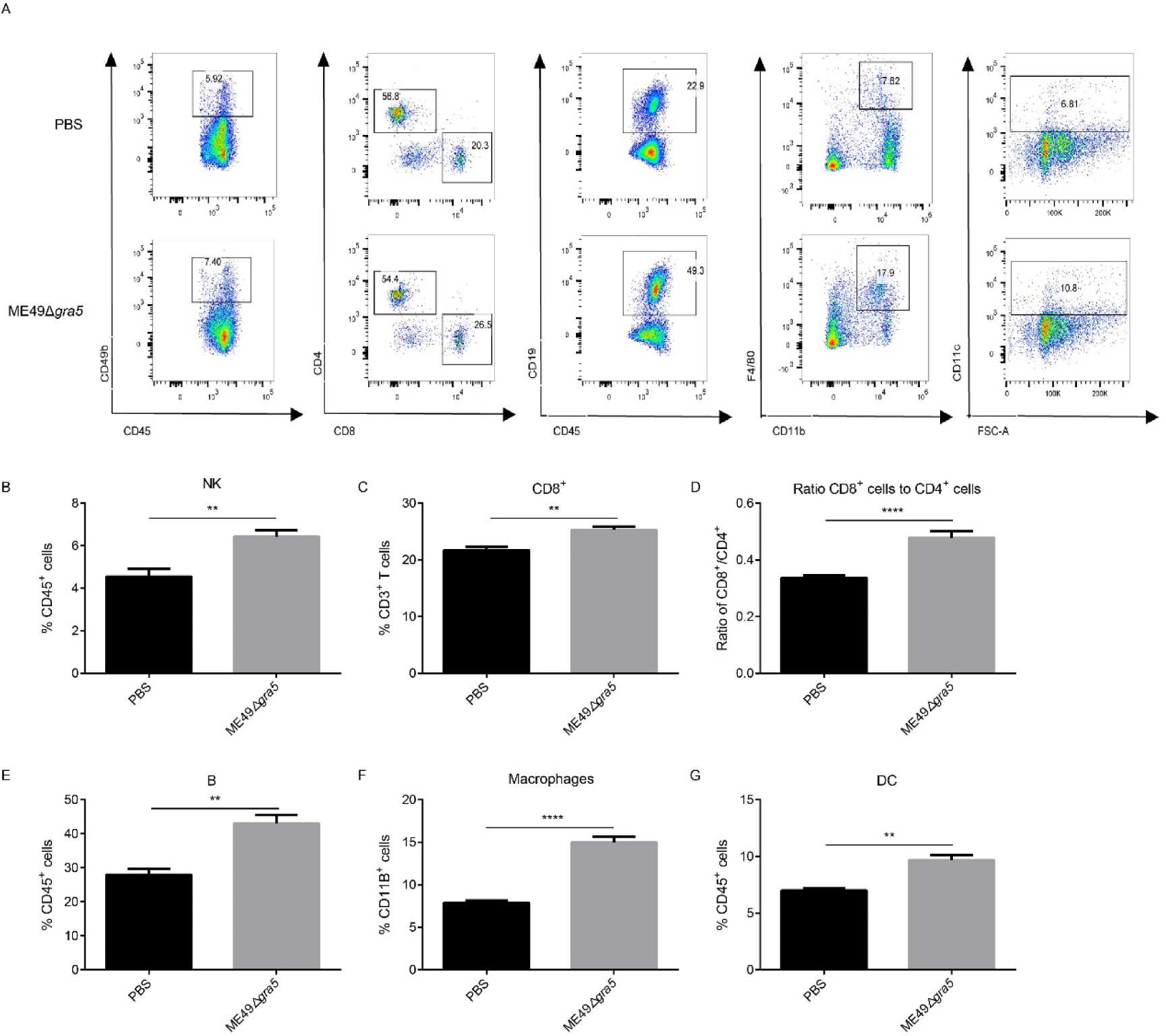
ME49Δ*gra5* therapy increased the percentages of innate and adaptive immune cells in the spleen. The mice bearing 4T1 tumors were inoculated with ME49-Δ*gra5* or PBS. The spleens of all mice were collected on day 21 after inoculation and the splenocyte was detected by flow cytometry. (A) Representative plots of flow cytometry data are presented in the panel. (B-G) The percentage of immune cells: NK cells (B), CD8^+^ T cells (C), B cells (E), macrophages (F), and DC cells (G); and the ratio of CD8^+^/CD4^+^ (D). Data are represented as mean ± SEM of three independent experiments (n=6). Statistical significance was assessed with a two-tailed unpaired Student’s *t*-test. \*\**p* <0.01, \*\*\*\**p* < 0.0001.

### ME49Δ*gra5* reduces the burden of the non-injected distant tumors

To determine whether ME49Δ*gra5* can activate systemic antitumor immunity, we constructed a bilateral 4T1 tumor mouse model. 4T1 cells were injected into both flanks of mice, followed by inoculating ME49Δ*gra5* tachyzoites *in situ* to the right flank tumor on day 9, 11, and 13 post 4T1 cell inoculation (Figure 8A). We found that ME49Δ*gra5* treatment inhibited the growth of the injected (right flank) and the distant (left flank) tumors (Figure 8B, C, D, E). Consequently, the survival of 4T1 bearing mice in ME49Δ*gra5* therapy group was much longer than that in the PBS treatment group (Figure 8F). The lung sections were stained with H&E. The results revealed fewer infiltrating myeloid cells and larger alveoli spaces in the ME49Δ*gra5* treatment group than that in the PBS treatment group (Figure 8G). Staining of the lung tissue with Ki67 to detect the 4T1 metastasis by immunohistochemistry (IHC) revealed a weaker lung metastasis with lower expression of Ki67 in ME49Δ*gra5* inoculated mice than that in the PBS treatment mice (Figure 8H, I).

**Figure 8.**
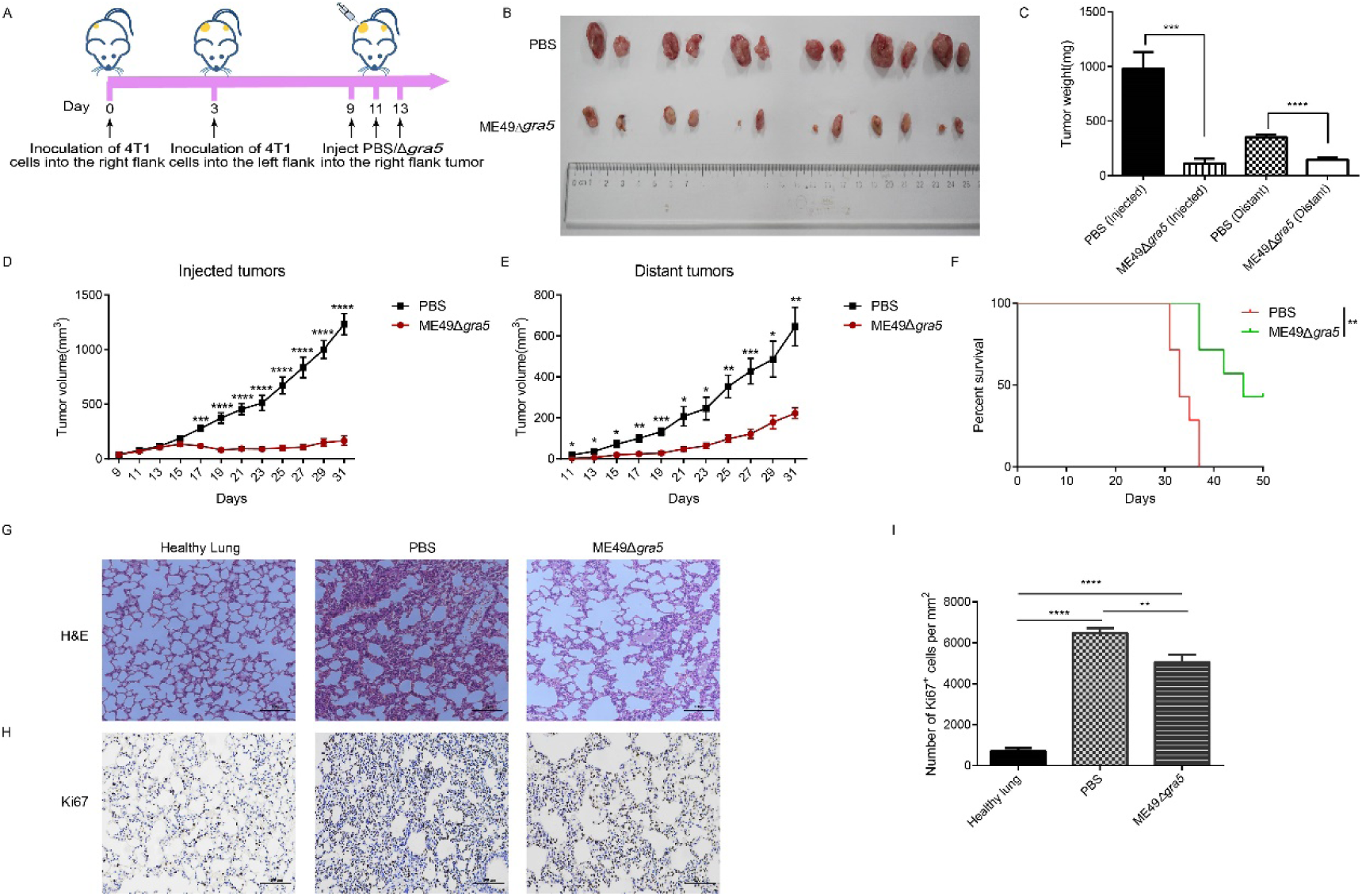
ME49Δ*gra5* parasites therapy suppressed the bilateral 4T1 tumor growth. (A)Schematic diagram to show the establishment processes of bilateral 4T1 mice model and treatment with ME49Δ*gra5* or PBS. (B-I) Comparison of the bilateral 4T1 tumor bearing mice treated with ME49Δ*gra5* or PBS: the bilateral 4T1 tumors (B); size of bilateral 4T1 tumors (C); volumes of the injected tumors (D); volumes of the distant tumors (E); survival curves (F); Lung sections stained with H&E (G); Immunohistochemistry staining of lung tissue with Ki67 antibody (H); the number of Ki67-positive cells in lung tissues (I). Data are represented as mean ± SEM of three independent experiments (A-F, n=7, G-I, n=3). Statistical significance was assessed with a two-tailed unpaired Student’s *t*-test (D, E), log-rank (Mantel–Cox) test (F), or One-way ANOVA (I). \**p*<0.05, \*\**p*<0.01, \*\*\**p*<0.001, \*\*\*\**p* <0.0001. Scale bar=100 μm.

To examine the immune system response and the immune status of the distant tumors, we first identified the cytokines that may suppress tumor growth. Mice serum, the parasite inoculated tumor and the distant tumor were collected, homogenized, and subjected to cytokines detection with ELISA. The levels of IFN-γ and IL12 were increased both in the parasite inoculated tumor and the distant tumor of ME49Δ*gra5* treated mice, and had no significant difference between these two TMEs (Figure S7A, B). The serum IFN-γ and IL12 showed the same increment in ME49Δ*gra5* therapy and the PBS treatment mice (Figure S7C, D). In addition, we detected the immune cells in the parasite inoculated and the distant tumors. The population of the innate immune cells and T cells, such as CD3+ (Figure 9A, B), CD4+ (Figure 9C, D), and CD8+ (Figure 9E, F), was significantly increased in the TMEs of ME49Δ*gra5* injected mice, compared to that of the PBS treatment mice.

**Figure 9.**
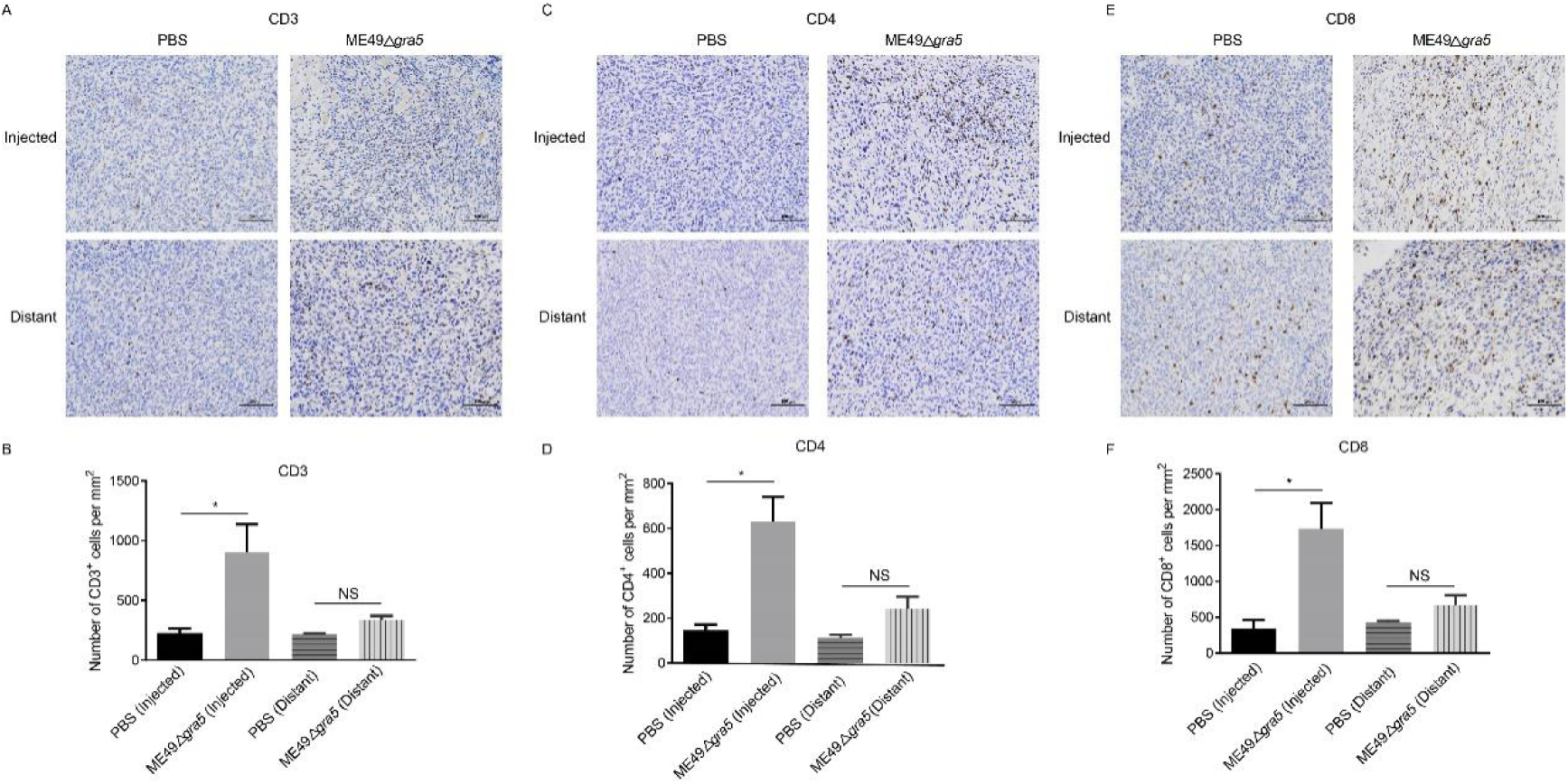
Detection of the immune status of the ME49-Δ*gra5* injected and distant tumors. (A-F) The bilateral 4T1 tumors were harvested on day 23 post 4T1 cell inoculation and used for immunohistochemistry staining for CD3 (A), CD4 (C), and CD8 (E); the numbers of CD3+ (B), CD4+ (D), and CD8+ (F) cells were calculated and compared between the PBS and the ME49-Δ*gra5* inoculation groups, for the injected and distant tumors. Data are represented as mean ± SEM of three independent experiments (n=3). Statistical significance was assessed with a two-tailed unpaired Student’s *t*-test. \**p*<0.05. Scale bar=100 μm.

## Discussion

GRA proteins are key factors in regulating cyst formation and maintenance, and so far, more that 60 GRA proteins have been identified [41]. The transmembrane domain-containing GRA proteins including GRA5 are translocated across the endoplasmic reticulum (ER) membrane into the lumen and exported from the ER, then transported from the Golgi to dense granules, and ultimately secreted out of the parasite [42, 43]. During the early differentiation of tachyzoite-to-bradyzoite transition, the intro-vacuole network (IVN)-associated GRA proteins including GRA1/2/4/6/9/12 translocate from the IVN to the cyst wall [2, 4]. Many bradyzoite specific GRAs have were identified, such as MCP3 knockouts form smaller cysts without other obvious defects [4], CST2 knockouts show an attenuated virulence and no cyst is detected in mice [4], GRA55 knockouts exhibits lower cyst burden in mice [44]. And BPK1 knockouts form smaller cysts more susceptible to pepsin digestion with reduced oral infection efficiency [45]. The function of many GRA proteins is poorly understood, particularly in the bradyzoite stage, and undoubtedly more GRA proteins are yet to be discovered [41]. In our research, in the brain homogenate of the mice infected by ME49Δ*gra5*, no cyst was detected at 75 dpi (Figure 1D); BAG1 protein band was detected weaker and weaker by WB at 15 and 30 dpi, and became undetectable at 75 dpi (Figure 1F); but the B1 gene which is present in both tachyzoites and bradyzoites replicated significantly at 15 dpi, compared to which at 7 dpi and 30 dpi, and then became undetectable at 75 dpi (Figure 1G). It may be due to the reason that, though ME49Δ*gra5* can be transformed into bradyzoites, deficiency of the cyst wall protein GRA5 resulted in defect of cyst wall formation, Gradually, the bradyzoite of *T. gondii* was eradicated by the host immunity.

Sequencing of the mice spleen transcriptome also presented an evidence of the activated host immunity against *T. gondii*, which revealed that most of the DEGs on immune related pathways in mice infected with ME49Δ*gra5* strain were down-regulated, compared to which in the mice infected with ME49-WT. For example, the DEGs involved in IL-17 signaling pathway or TNF signaling pathway were down-regulated, including Mmp9, IL1b, Cxcl5, Cxcl1, Ccl7, Ccl2, Mmp3, Creb5 and other genes. IL-17 is known as an early initiator of T cell induced inflammatory response, which can amplify inflammatory response by promoting the release of proinflammatory cytokines, downregulation of IL-17 signaling pathway helps prevent the fatal inflammation caused by *T. gondii* infection [46]. IFN-γ plays a main role in immune protection but also brings immunopathologic injury to hosts when dysregulated [47]. IL-10 knockout mice infected with *T. gondii* succumb to a lethal immune response of overproduced IL12 and IFN-γ [48]. In this study, the survival rate of ME49Δ*gra5* infected mice was considerably higher than that of ME49-WT infected mice. The reason may be that ME49Δ*gra5* induced a downregulated transcription of the genes involved in IL-17 signaling pathway and excessive inflammatory reaction, which stimulated immune responses to maintain immune balance.

Anti-*T. gondii* vaccines mainly include inactivated vaccine, nucleic acid vaccine, recombinant subunit vaccine, and attenuated live vaccine. The inactivated vaccine is prepared by collecting dead *T. gondii* tachyzoites [49]. Nucleic acid vaccine is developed by introducing multiple genes or a single gene coding the protein capable of eliciting immune responses, into overexpression plasmids or viral vectors. The research on *T. gondii* nucleic acid vaccines has mainly focused on the gene encoding virulence factors or host invasion-related proteins, such as rhoptry proteins [50, 51], dense granule proteins [52, 53], and microneme proteins [54]. Recombinant subunit vaccine is prepared by expressing parasite proteins or motifs with strong immunogenicity, such as GRA2 and GRA5 [37]. Although these vaccines are safe, they are not potent vaccines due to their relatively short maintenance or poor effects of immune protection. Compared with other vaccines, the attenuated life vaccines can accurately simulate the natural infection of *T. gondii*, which may induce more effective immune protection. The attenuated α-*amy* or *gra9* knockout strains can serve as vaccines against acute and chronic *T. gondii* infection [55, 56]. However, as a few cysts are still formed by these attenuated strains, the safety is challenged. In this study, no cyst was observed being formed by the ME49Δ*gra5* mutant, meanwhile it induced a long-term immune response in mice. Several important cytokines, including IFN-γ, IL12, and TNF-α, were induced to secret at high levels in Balb/c mice after vaccination with ME49Δ*gra5* tachyzoites. According to the genotype and the virulence in murine model, *T. gondii* is mainly divided into type I, type II, and type III strains, such as RH, ME49 and VEG strains, respectively [57]. These strains have different proliferation abilities and pathogenicity in humans and animals, which will bring challenges to the prevention and control of toxoplasmosis. Here, ME49Δ*gra5* immunization to mice achieved 100% protection against challenges with tachyzoites of these three strains and ME49 cysts. Therefore, ME49Δ*gra5* vaccination is effective against a broad spectrum of *T. gondii* strains.

The immune response is involved in the occurrence and development of tumors, including non-specific immunity and specific immunity, of which specific immunity is divided into cellular immunity and humoral immunity. The differentiated macrophages have two phenotypes that one is classically activated macrophages (M1) and the other is alternatively activated macrophages (M2) [58]. M1 macrophages can be activated by IFN-γ to produce pro-inflammatory, which is good for anti-tumor, while M2 macrophages promote the proliferation, invasion, and metastasis of tumor cells [59]. NK cells are cytotoxic innate lymphoid cells (ILCs), which are able to eliminate malignant cells and limit tumor metastases through secreting IFN-γ [60]. Cellular immunity is the main way of the immune responses against tumors, including CD4^+^ T cells and CD8^+^ T cells [61]. Cytotoxic CD8^+^ T cells, activated by antigen-presenting cells, are the most vital in anti-cancer immune responses and constitute the backbone of cancer immunotherapy [62]. CD4^+^ T cells can directly kill the tumor cells by producing a high level of IFN-γ, moreover, it can enhance anti-tumor cytotoxic T-lymphocytes (CTLs) response by promoting clonal replication at the tumor site and acting as APC for CTLs to preferentially generate immune memory cells [63]. In addition to IFN-γ, IL12 also plays an essential role in anti-tumor immunity. IL12 is beneficial to cytotoxic lymphocyte maturation and is a stimulatory factor of NK cells [64, 65]. The levels of IFN-γ and IL12 was much higher both in TME and serum of the 4T1 bearing mice immunized with ME49Δ*gra5* tachyzoites. Furthermore, ME49Δ*gra5 in situ* vaccination also elevated the levels of IFN-γ and IL12 in distant TME. The immune cells that process and present tumor antigens, such as DCs, and the cells killing tumors, such as NK cells, CD4^+^ T cells, and CD8^+^ T cells, were all significantly increased in the spleens of ME49Δ*gra5* vaccinated mice. Further, the number of infiltrated CD3^+^, CD4+ and CD8^+^ T cells also significantly elevated in the TME with ME49Δ*gra5* parasites injection compared to which of the TME with PBS injection, but no significant difference was found with these infiltrated cells in the TME of the distant 4T1 tumor and in the TME with PBS infection. These results indicated that these immune cells have an important role in restraining tumor growth at the site with ME49Δ*gra5* infection. The antibodies and cytokines induced by this parasite infection are key factors in the inhibition of *T. gondii* tachyzoites and tumor cells. Intratumoral injection of the avirulent Δ*gra17* mutant showed similar mechanism [36]. However, Δ*gra17* mutant formed few brain cysts in infected mice [66]. In our research, ME49Δ*gra5* parasites can not form brain cysts, which implied it may be safer than Δ*gra17* mutant. Taken together, ME49Δ*gra5* parasites inhibited tumor growth through immune modulation.

Host cells produce IL-12 and IFN-γ to resist *T. gondii* infection, which is an important immune protection mechanism of the host against *T. gondii* infection [67, 68]. Dendritic cells, macrophages, NK cells, T lymphocytes, and a variety of cytokines are involved in the responses to *T. gondii* infection [69, 70]. Macrophages and dendritic cells recognize parasitic antigens and promote the immune cells to produce IL12, which activates and stimulates the proliferation of NK cells, CD4^+^ T cells, and CD8^+^ T cells [71–73]. In addition, IL-12 activates T cells and NK cells to produce IFN-γ [67]. The host anti-*T. gondii* immune response exhibits many similarities with anti-tumor response. More and more studies are focused on the application of *T. gondii* as a cancer immunotherapeutic agent. Among them uracil auxotrophic mutants of *T. gondii* was effective in treatment of melanoma, ovarian cancer, and pancreatic cancer [74, 75]. In our study, we demonstrated that 4T1 intratumor injection with ME49Δ*gra5,* suppressed the growth of both the injected tumor and the non-injected distant tumor, and prevented 4T1’s lung metastasis. Combination therapy shows more efficiency in anti-tumor. For example, when mice were treated with a combination of CPMV and CPA, the 4T1 tumor volume was much smaller than that of the mice treated with CPMV alone [31]. Combination therapy of ME49Δ*gra5* need further study.

In summary, as shown in Figure 10, our results demonstrated that GRA5 is critical for *T. gondii*’s virulence and cyst formation. ME49Δ*gra5* tachyzoites vaccination triggered the generation of pro-inflammatory factors to protect mice from challenge infection of three types of *T. gondii,* RH, ME49 and VEG, and this protection was long-lasting. Besides, intratumoral injection of ME49Δ*gra5* showed a high efficacy against the growth of the injected and distant 4T1 tumors, as well as lung metastasis. ME49Δ*gra5* inoculation promoted the production of IFN-γ and IL12, increased the innate and adaptive immune cells of the spleen and tumor-infiltrating. ME49Δ*gra5* was shown to be a promising vaccine against *T. gondii* infection and a potential immunotherapeutic agent against tumors.

**Figure 10.**
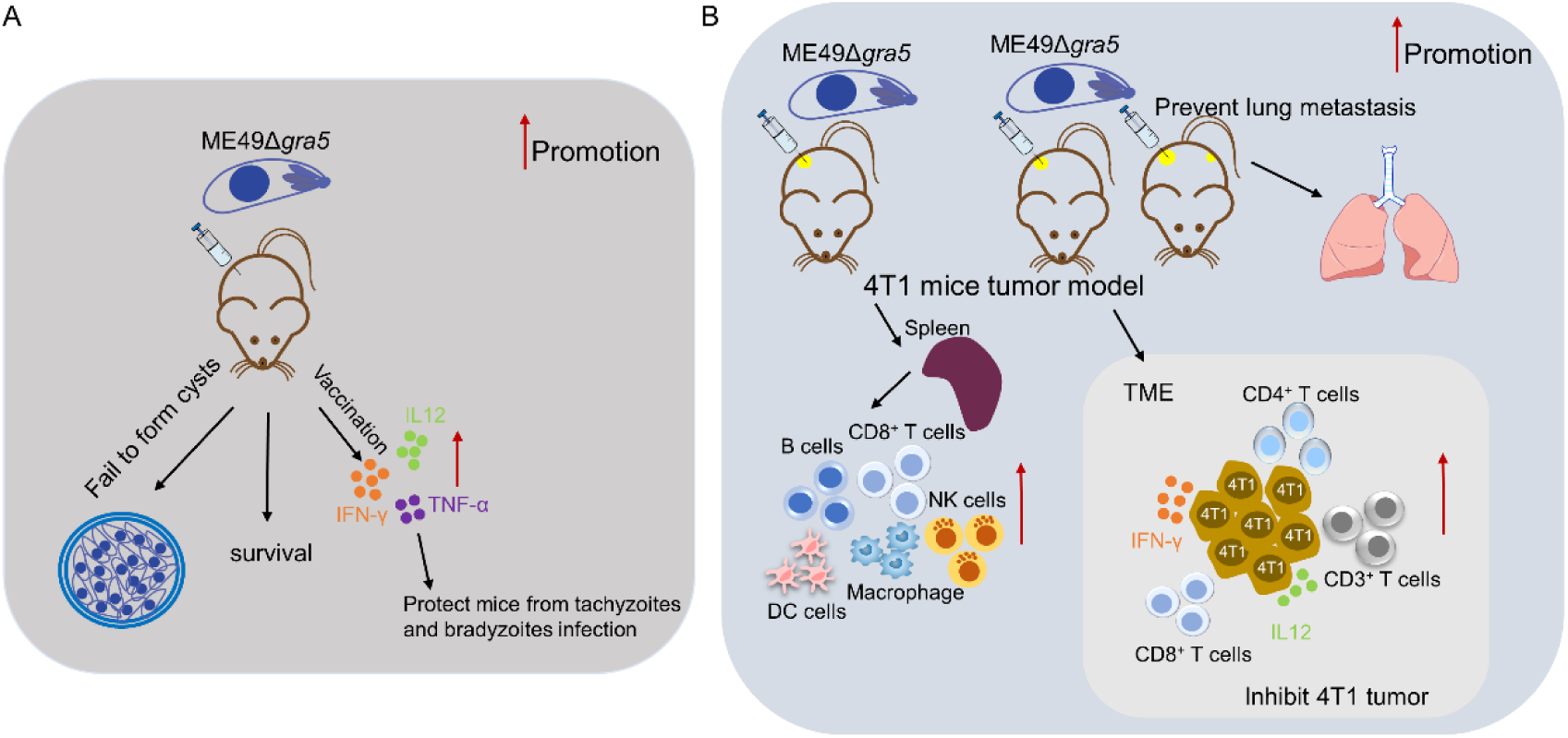
Summary for the potential of ME49Δ*gra5* as an attenuated live vaccine against *T. gondii* infection and immuno-therapeutic agent against 4T1 tumor. (A) ME49Δ*gra5* vaccination against *T. gondii* infection; (B) ME49Δ*gra5* inoculation against 4T1 tumor.

## Materials and methods

### Animal and animal ethics

Specific pathogen free (SPF) female Balb/c mice of 6-8 weeks old, and SPF male SV129 mice of 8 weeks old, were purchased from the Laboratory Animal Centre and raised in an SPF laboratory in the university. All animal experiments were performed following the guidelines for laboratory animal manipulation.

### Parasites and cells

ME49-WT and ME49Δ*gra5* parasites were maintained in human foreskin fibroblasts (HFF, American Type Culture Collection, ATCC, USA) cells. HFF cells were cultured in Dulbecco’s modified Eagle’s medium (DMEM, Gibco, USA) and 4T1 cells were cultured in Roswell Park Memorial Institute-1640 (RPMI-1640, Gibco) supplemented with 10% fetal bovine serum (FBS, Gibco) and 1% Penicillin-Streptomycin (Thermo Fisher Scientific) at 37°C and 5% CO_2_.

### Antibodies

The antibodies used for western blot: rabbit monoclonal anti-β-actin was purchased from Cell Signaling Technology, and mouse monoclonal anti-BAG1 was raised in our lab. immunofluorescence assay: mouse monoclonal anti-SAG1, goat anti-mouse IgG H&L (Alexa Fluor® 488), and goat anti-mouse IgG H&L (Alexa Fluor® 594), were purchased from Abcam (United Kingdom). The antibodies used for Immunohistochemistry: anti-Ki67 rabbit polyclonal antibody, anti-CD3 rabbit polyclonal antibody, anti-CD4 rabbit polyclonal antibody, and anti-CD8 rabbit polyclonal antibody, were purchased from Servicebio (China).

### Construction of *T. gondii* ME49Δ*gra5* strain

The *gra5* knockout ME49 strain (ME49Δ*gra5*) was constructed by CRISPR-Cas9 technology (Figure S1 in supplemental data). The homologous template and the sgRNA CRISPR plasmid were co-transfected to ME49-WT parasites, and the parasites were screened with 3 μM pyrimethamine (Sigma, Germany). The stable lines of gene knockout parasites were confirmed by PCR. The sequences of the primers used in generation were shown in Table S1.

### Immunofluorescence assay (IFA)

For invasion assay, the cells were fixed with 4% paraformaldehyde for 5 min at room temperature (RT) and then blocked with 10% bovine serum albumin (BSA) in PBS for 1 h at 37°C, followed by incubation in the mouse anti-*Tg*SAG1 monoclonal primary antibody (Abcam) (1:800 diluted in PBS) at 4°C overnight. The cells were then washed for 3 times with PBS, and after that, incubated with the Alexa Fluor® 488 conjugated goat anti-mouse IgG secondary antibody (Invitrogen, United States) (1:2000 diluted in PBS) for 1 h at 37°C. Following, the cells were permeabilized with 0.5% Triton X-100 (DingGuo, China) for 10 min at RT and blocked with 10% BSA in PBS for 1 h at 37°C. The cells were incubated again with the mouse anti-*Tg*SAG1 monoclonal primary antibody (Abcam) at 4°C overnight, followed by incubation with the Alexa Fluor® 594 conjugated goat anti-mouse IgG secondary antibody (Invitrogen) for 1 h at 37°C. After washing for 3 times with PBS, the coverslips were taken out and rinsed with double distilled water (ddH_2_O), and mounted with DAPI mounting oil (Southern Biotech, United States).

For proliferation assay, the cells were fixed with 4% paraformaldehyde for 10 min at RT, and then permeabilized with 0.5% Triton X-100 for 10 min at RT and blocked with 10% BSA in PBS for 1 h at 37°C. The cells were incubated with the mouse anti-*Tg*SAG1 monoclonal primary antibody (Abcam) at 4°C overnight, followed by incubation with the Alexa Fluor® 488 conjugated goat anti-mouse IgG secondary antibody (Invitrogen) for 1 h at 37°C. After that, the coverslips were washed with PBS, rinsed with ddH_2_O and mounted with DAPI mounting oil (Southern Biotech).

### Mice spleen transcriptome sequencing

The female Balb/c mice of 6-8 weeks old were infected with 10^3^ freshly egressed tachyzoites of ME49-WT or ME49Δ*gra5* by i.p. injection, or injected a same volume of PBS for control, 3 mice for each group. At 7 dpi, mice were sacrificed and dissected. The spleens were collected and subjected to total RNA extraction with TRIzol^TM^ Agent (Invitrogen). The Total RNA was sent to Shanghai Major Biology (China) for transcriptome sequencing.

### Comparison of the acute virulence (parasite’s invasion, proliferation and host survival) for ME49 and ME49Δ*gra5*

HFF cells were seeded in a 12-well plates, grew to 100% confluent, and infected with ME49-WT or ME49Δ*gra5* strains (3 wells for each group) at multiplicity of infection (MOI) 1 for 1 h, and then washed with PBS for 3 times. For invasion assay, the cells were subjected to IFA immediately. For proliferation assay, the cells were further cultured for 24 h, and then subjected to IFA after washing 3 times with PBS. The number of parasites in each parasitophorous vacuole (PV), and the average number of parasites per PV were calculated.

For survival assay, 6-8 weeks old Balb/c female mice were i.p. injected with 10^3^ ME49-WT tachyzoites, or 10^3^, 10^4^, 10^5^, and 10^6^ ME49Δ*gra5* tachyzoites respectively, and the survival of mice was monitored for 30 days (n=6 mice). The sera of the infected mice were collected from tails at 7 dpi for ME49-WT infection, or at 7, 14, 21, 30 dpi for ME49Δ*gra5* infection, and subjected to antibody detection by MAT [76]. The sera of the ME49-WT chronically infected mice were used as the positive control.

### Cysts formation assay for ME49-WT and ME49Δ*gra5*

The SV129 male mice of 8-week-old were i.p. challenged with 100 fresh tachyzoites of ME49-WT or ME49Δ*gra5* (3 mice per group). After 75 dpi, the mice were sacrificed, and the brains were harvested and homogenized, and fixed. The brain homogenate was smeared on slides and stained with FITC-conjugated Dolichos Biflorus Agglutinin (DBA) antibody (Vector Laboratories, USA), and then cysts were calculated under a fluorescence microscope. The brain homogenate was subjected to B1 gene copy detection with q-PCR, the primers are provided in Table S1.

We further analyzed the cyst formation for ME49Δ*gra5* at different time post infection. Balb/c female mice of 6-8 weeks old were i.p. injected with 10^3^ ME49Δ*gra5* tachyzoites, and the mice i.p. infected with 100 ME49-WT tachyzoites for 75 days were used as the positive control (n=3). The mice were sacrificed, and the brains were harvested and homogenized. The BAG1 protein in each brain homogenates were detected by western blot at 15, 30, and 75 dpi; the B1 copies in each brain homogenate was detected by qPCR at 7, 15, 30, 75, and 150 dpi.

### Evaluation of the short and long term protective efficiency for ME49Δ*gra5* vaccination

For protective efficiency assay, 6-8 weeks old female Balb/c mice were first vaccinated with 10^3^ freshly egressed tachyzoites of ME49Δ*gra5* for 30 or 75 days. On day 30 and 75 post vaccination, the tail blood of immunized mice and naïve mice were obtained for detection, and then these mice were i.p. injected with 10^3^ freshly egressed tachyzoites of RH-WT, ME49-WT, or VEG-WT (8 mice per group). Moreover, another two groups of immunized mice and naïve mice were orally infected with 20 fresh brain cysts of ME49 to assess the efficacy against bradyzoites infection (5 mice per group). These mice were monitored daily for another 30 days.

### Evaluation of the parasitic burden *in vivo*

The female Balb/c mice of 6-8 weeks old were infected with 10^3^ freshly egressed tachyzoites of ME49-WT or ME49Δ*gra5* by i.p. injection (5 mice per group). Before infection and 7 dpi, each mouse was weighted and the weight change was recorded. At 7 dpi, mice were sacrificed and dissected. The murine brains, livers, spleen, and lungs were collected and weighed respectively. Ten milligram spleen, 25 mg brain, lung, liver, and 1 ml peritoneal fluid, were collected, and used for genomic DNA extraction with DNeasy Blood & Tissue Kit (Qiagen, Germany). The parasitic burden from each sample was determined by detecting the B1 gene copies with qPCR using Hieff® qPCR SYBR® Green Master Mix (Low Rox Plus) (Yeasen, China).

### Construction mouse tumor model and treatment of mice with ME49Δ*gra5*

The 4T1 murine breast tumor cells were cultured and harvested, and then resuspended in RPMI-1640 medium at a concentration of 1 × 10^6^ cells per ml. For the unilateral 4T1 mice tumor model, 5-6 weeks old Balb/c female mice were subcutaneously (s.c.) injected on the right flank with 100 μl of the 4T1 cell mixture (10^5^ cells per mouse), 7 mice per group. Tumors were allowed to grow for 9 days, and then 50 μl PBS or 1 × 10^5^ ME49Δ*gra5* parasites were intratumorally injected on day 9, 11, and 13 post 4T1 cell injection. Following, the tumor size in each mouse was recorded every 2 days, and mice survival was also monitored. Until 21 dpi, the mice were sacrificed, and the tumors were taken out and photographed. For the bilateral tumor model, mice were s.c. injected on the right flank with 100 μl of the 4T1 cell mixture (10^5^ cells per mouse), 7 mice per group. Three days later, mice were s.c. inoculated on the left flank with the same amount of 4T1 cells. After 3 days (tumors were allowed to grow for 6 days on the left flank, and 9 days on the right flank), 50 μl PBS or 1 × 10^5^ ME49Δ*gra5* parasites in 50 ul PBS were injected into the right flank tumor. Tumor size and mice survival were measured every 2 days. Until 23 dpi, the mice were sacrificed, and the tumors were taken out and photographed.

### Cytokine assay

Mice serum and tumor tissue were collected to detect the level of IFN-γ and IL12p40 with ELISA kits (Multi Sciences, China) based on the manufacturer’s recommendations. Mice serum was collected from the naïve mice or ME49Δ*gra5* immunized mice at 30 or 75 dpi, and the tumor-bearing mice treated with PBS or ME49Δ*gra5* tachyzoites as described above. For tumor tissue, 25 mg of tumor tissue was weighed, and ground with 500 μl pre-cooled PBS to single-cell suspension. The suspension was stored at -80°C for 30 min, and then taken out, and dissolved completely in a water bath at 37°C. The suspension was mixed up and down, and placed back at -80°C. After the freezing-thaw was repeated for 3 times, the suspension was centrifuged at 4°C, 5000×g, for 5 min, and the supernatant was taken for ELISA detection.

### Flow cytometry

The mice were inoculated with 4T1 cells and were treated with ME49Δ*gra5* parasites or PBS as described above. At 21 dpi, six mice from each group were euthanized to harvest the spleens which were homogenized and filtered to get the spleen single cell suspension, and red blood cells were lysed with erythrocyte lysis buffer. The concentration of splenocytes was adjusted to 1 × 10^6^ cells/ml with PBS after washing with RPMI-1640 medium twice. And then, the single-cell suspensions were incubated with the antibodies (Univ, China) of panel 1: percp-cy5.5 anti-mouse CD45, BV510 anti-mouse CD3e, APC anti-mouse CD4, FITC anti-mouse CD8a, PE-cy7 anti-mouse CD19, PE anti-mouse CD49b, panel 2: percp-cy5.5 anti-mouse CD45, PE anti-mouse F4/80, FITC anti-mouse CD11b and APC anti-mouse CD11c, at 4°C for 30 min following the manufacturer’s guidance. The analyses of splenocytes were performed with fluorescence profiles by a FACScan flow cytometer (BD Biosciences, USA).

### Hematoxylin and eosin (H&E) staining

The lungs were cut into 4 μm sections. The sections were dewaxing and stained with Hematoxylin solution for 3-5 min, followed by 85% ethanol, and 95% ethanol dehydration. Finally, the sections were stained with Eosin dye for 5 min and dehydrated. The samples were observed, and photos were captured under a light microscope, with nucleus stained in blue and cytoplasm stained in red.

### Immunohistochemistry (IHC)

The lungs or tumors were cut into 4 μm sections. The sections were dewaxed in xylene I and II for 15 min, and dehydrated in (100%, 85%, and 75%) alcohol for 5 min each, and rinsed with ddH_2_O. The samples were placed in citric acid (pH6.0) for antigen repair. After that, the sections were put into 3% hydrogen peroxide for 25 min at RT in the dark and blocked with 3% BSA in PBS for 30 min at RT, followed by incubating with the primary antibodies (Servicebio) at 4°C overnight. Following, the slides were washed with BPS for 5 min with gentle shaking for 3 times, and then incubated with the goat anti-rabbit IgG H&L (HRP) antibody for 50 min at RT. DAB (3,3-N-Diaminobenzidine Tertrahydrochloride) was used for staining and hematoxylin was used for counter-staining of nuclei. Then, the sections were dehydrated and mounted for observation and photos capture under a light microscope, with nucleus stained blue and DAB-positive stained brownish yellow.

### Statistical Analysis

SPSS20 statistical software was used to analyze the data. The analysis of the difference between two groups were conducted by unpaired 2-tailed Student’s *t*-test. One-way ANOVA or Kruskal-Wallis test was performed to compare the difference among more than two groups. The survival of mice was compared using log-rank (Mantel–Cox) test. The statistical significance was defined as *p*<0.05*.

## Supplementary information

This article contains 1 supplemental table and 7 supplemental figures.

## Author’s contributions

MC and PY performed the experiments, analyzed data and wrote the manuscript. ZXX performed qPCR assays. JTC, WHZ performed WB assay. LJZ, LLY and JP performed flow cytometry. HP performed study conception and designing, supervision of the research group, funding support, and drafting the manuscript. All authors read and approved the final version of the manuscript.

## Acknowledgement

We greatly appreciate Professor Dominique Soldati-Favre from University of Geneva for kind critiques and suggestions to our research.

## Funding supports

This research was supported by National Natural Science Foundation of China (82272364, 81971954), Key project of Guangzhou science research (201904020011), and the Basic Research Project of Key Laboratory of Guangzhou (202102100001) to HJP. National Natural Science Youth Foundation of China (32102693), Medical Science and Technology Foundation of GuangDong Province (A2021167) to LJZ.

## Data availability

The datasets supporting the findings of this research are included within the article.

## References

1. Tomita T, Mukhopadhyay D, Han B, Yakubu R, Tu V, Mayoral J, et al. Toxoplasma gondii Matrix Antigen 1 Is a Secreted Immunomodulatory Effector. mBio. 2021;12(3). doi: 10.1128/mBio.00603-21. PubMed PMID: 34006649; PubMed Central PMCID: PMC8262993.

2. Guevara RB, Fox BA, Falla A, Bzik DJ. Toxoplasma gondii Intravacuolar-Network-Associated Dense Granule Proteins Regulate Maturation of the Cyst Matrix and Cyst Wall. mSphere. 2019;4(5). Epub 2019/10/18. doi: 10.1128/mSphere.00487-19. PubMed PMID: 31619500; PubMed Central PMCID: PMCPMC6796980.

3. Sibley LD. Intracellular parasite invasion strategies. Science. 2004;304(5668):248–53. doi: 10.1126/science.1094717. PubMed PMID: 15073368.

4. Tu V, Mayoral J, Sugi T, Tomita T, Han B, Ma YF, et al. Enrichment and Proteomic Characterization of the Cyst Wall from In Vitro Toxoplasma gondii Cysts. mBio. 2019;10(2). Epub 2019/05/02. doi: 10.1128/mBio.00469-19. PubMed PMID: 31040239; PubMed Central PMCID: PMCPMC6495374.

5. Mercier C, Cesbron-Delauw MF. Toxoplasma secretory granules: one population or more? Trends in parasitology. 2015;31(11):604. doi: 10.1016/j.pt.2015.02.002. PubMed PMID: 29496113.

6. Torpier G, Charif H, Darcy F, Liu J, Darde ML, Capron A. Toxoplasma gondii: differential location of antigens secreted from encysted bradyzoites. Experimental parasitology. 1993;77(1):13–22. doi: 10.1006/expr.1993.1056. PubMed PMID: 8344403.

7. Hill DE, Chirukandoth S, Dubey JP. Biology and epidemiology of Toxoplasma gondii in man and animals. Animal health research reviews. 2005;6(1):41–61. doi: 10.1079/ahr2005100. PubMed PMID: 16164008.

8. Chen M, Zhou L, Li S, Wei H, Chen J, Yang P, et al. Toxoplasma gondii DNA methyltransferases regulate parasitic energy metabolism. Acta tropica. 2022;229:106329. Epub 2022/02/06. doi: 10.1016/j.actatropica.2022.106329. PubMed PMID: 35122712.

9. Sugi T, Tu V, Ma Y, Tomita T, Weiss LM. Toxoplasma gondii Requires Glycogen Phosphorylase for Balancing Amylopectin Storage and for Efficient Production of Brain Cysts. mBio. 2017;8(4). Epub 2017/08/31. doi: 10.1128/mBio.01289-17. PubMed PMID: 28851850; PubMed Central PMCID: PMCPMC5574715.

10. Vesely MD, Zhang T, Chen L. Resistance Mechanisms to Anti-PD Cancer Immunotherapy. Annual review of immunology. 2022;40:45–74. Epub 2022/04/27. doi: 10.1146/annurev-immunol-070621-030155. PubMed PMID: 35471840.

11. Bruni D, Angell HK, Galon J. The immune contexture and Immunoscore in cancer prognosis and therapeutic efficacy. Nature reviews Cancer. 2020;20(11):662–80. doi: 10.1038/s41568-020-0285-7. PubMed PMID: 32753728.

12. Fridman WH, Pages F, Sautes-Fridman C, Galon J. The immune contexture in human tumours: impact on clinical outcome. Nature reviews Cancer. 2012;12(4):298–306. doi: 10.1038/nrc3245. PubMed PMID: 22419253.

13. Reuschenbach M, von Knebel Doeberitz M, Wentzensen N. A systematic review of humoral immune responses against tumor antigens. Cancer immunology, immunotherapy : CII. 2009;58(10):1535–44. doi: 10.1007/s00262-009-0733-4. PubMed PMID: 19562338; PubMed Central PMCID: PMC2782676.

14. Shan L, Flavell RA, Herndler-Brandstetter D. Development of Humanized Mouse Models for Studying Human NK Cells in Health and Disease. Methods in molecular biology (Clifton, NJ). 2022;2463:53–66. Epub 2022/03/29. doi: 10.1007/978-1-0716-2160-8_5. PubMed PMID: 35344167; PubMed Central PMCID: PMCPMC9116980.

15. Hinshaw DC, Shevde LA. The Tumor Microenvironment Innately Modulates Cancer Progression. Cancer research. 2019;79(18):4557–66. doi: 10.1158/0008-5472.CAN-18-3962. PubMed PMID: 31350295; PubMed Central PMCID: PMC6744958.

16. Hu H, Chen Y, Tan S, Wu S, Huang Y, Fu S, et al. The Research Progress of Antiangiogenic Therapy, Immune Therapy and Tumor Microenvironment. Frontiers in immunology. 2022;13:802846. doi: 10.3389/fimmu.2022.802846. PubMed PMID: 35281003; PubMed Central PMCID: PMC8905241.

17. Ferlay J, Colombet M, Soerjomataram I, Mathers C, Parkin DM, Pineros M, et al. Estimating the global cancer incidence and mortality in 2018: GLOBOCAN sources and methods. Int J Cancer. 2019;144(8):1941–53. Epub 2018/10/24. doi: 10.1002/ijc.31937. PubMed PMID: 30350310.

18. Parsons J, Francavilla C. ’Omics Approaches to Explore the Breast Cancer Landscape. Frontiers in cell and developmental biology. 2019;7:395. doi: 10.3389/fcell.2019.00395. PubMed PMID: 32039208; PubMed Central PMCID: PMC6987401.

19. Rossi C, Cicalini I, Cufaro MC, Consalvo A, Upadhyaya P, Sala G, et al. Breast cancer in the era of integrating “Omics” approaches. Oncogenesis. 2022;11(1):17. doi: 10.1038/s41389-022-00393-8. PubMed PMID: 35422484.

20. Vagia E, Mahalingam D, Cristofanilli M. The Landscape of Targeted Therapies in TNBC. Cancers (Basel). 2020;12(4). Epub 2020/04/12. doi: 10.3390/cancers12040916. PubMed PMID: 32276534; PubMed Central PMCID: PMCPMC7226210.

21. Cardoso F, Senkus E, Costa A, Papadopoulos E, Aapro M, Andre F, et al. 4th ESO-ESMO International Consensus Guidelines for Advanced Breast Cancer (ABC 4)dagger. Ann Oncol. 2018;29(8):1634–57. Epub 2018/07/23. doi: 10.1093/annonc/mdy192. PubMed PMID: 30032243; PubMed Central PMCID: PMCPMC7360146.

22. Waks AG, Winer EP. Breast Cancer Treatment: A Review. JAMA. 2019;321(3):288–300. Epub 2019/01/23. doi: 10.1001/jama.2018.19323. PubMed PMID: 30667505.

23. Kim C, Gao R, Sei E, Brandt R, Hartman J, Hatschek T, et al. Chemoresistance Evolution in Triple-Negative Breast Cancer Delineated by Single-Cell Sequencing. Cell. 2018;173(4):879–93 e13. Epub 2018/04/24. doi: 10.1016/j.cell.2018.03.041. PubMed PMID: 29681456; PubMed Central PMCID: PMCPMC6132060.

24. Lyon AR, Yousaf N, Battisti NML, Moslehi J, Larkin J. Immune checkpoint inhibitors and cardiovascular toxicity. Lancet Oncol. 2018;19(9):e447–e58. Epub 2018/09/08. doi: 10.1016/S1470-2045(18)30457-1. PubMed PMID: 30191849.

25. Sivan A, Corrales L, Hubert N, Williams JB, Aquino-Michaels K, Earley ZM, et al. Commensal Bifidobacterium promotes antitumor immunity and facilitates anti-PD-L1 efficacy. Science. 2015;350(6264):1084–9. doi: 10.1126/science.aac4255. PubMed PMID: 26541606; PubMed Central PMCID: PMC4873287.

26. Toso JF, Gill VJ, Hwu P, Marincola FM, Restifo NP, Schwartzentruber DJ, et al. Phase I study of the intravenous administration of attenuated Salmonella typhimurium to patients with metastatic melanoma. Journal of clinical oncology : official journal of the American Society of Clinical Oncology. 2002;20(1):142–52. doi: 10.1200/JCO.2002.20.1.142. PubMed PMID: 11773163; PubMed Central PMCID: PMC2064865.

27. Lizotte PH, Baird JR, Stevens CA, Lauer P, Green WR, Brockstedt DG, et al. Attenuated Listeria monocytogenes reprograms M2-polarized tumor-associated macrophages in ovarian cancer leading to iNOS-mediated tumor cell lysis. Oncoimmunology. 2014;3:e28926. doi: 10.4161/onci.28926. PubMed PMID: 25083323; PubMed Central PMCID: PMC4106169.

28. Chowdhury S, Castro S, Coker C, Hinchliffe TE, Arpaia N, Danino T. Programmable bacteria induce durable tumor regression and systemic antitumor immunity. Nature medicine. 2019;25(7):1057–63. doi: 10.1038/s41591-019-0498-z. PubMed PMID: 31270504; PubMed Central PMCID: PMC6688650.

29. Bommareddy PK, Shettigar M, Kaufman HL. Integrating oncolytic viruses in combination cancer immunotherapy. Nature reviews Immunology. 2018;18(8):498–513. doi: 10.1038/s41577-018-0014-6. PubMed PMID: 29743717.

30. Lizotte PH, Wen AM, Sheen MR, Fields J, Rojanasopondist P, Steinmetz NF, et al. In situ vaccination with cowpea mosaic virus nanoparticles suppresses metastatic cancer. Nature nanotechnology. 2016;11(3):295–303. doi: 10.1038/nnano.2015.292. PubMed PMID: 26689376; PubMed Central PMCID: PMC4777632.

31. Cai H, Wang C, Shukla S, Steinmetz NF. Cowpea Mosaic Virus Immunotherapy Combined with Cyclophosphamide Reduces Breast Cancer Tumor Burden and Inhibits Lung Metastasis. Advanced science. 2019;6(16):1802281. doi: 10.1002/advs.201802281. PubMed PMID: 31453050; PubMed Central PMCID: PMC6702650.

32. Kim JO, Jung SS, Kim SY, Kim TY, Shin DW, Lee JH, et al. Inhibition of Lewis lung carcinoma growth by Toxoplasma gondii through induction of Th1 immune responses and inhibition of angiogenesis. Journal of Korean medical science. 2007;22 Suppl:S38-46. doi: 10.3346/jkms.2007.22.S.S38. PubMed PMID: 17923753; PubMed Central PMCID: PMC2694397.

33. Baird JR, Byrne KT, Lizotte PH, Toraya-Brown S, Scarlett UK, Alexander MP, et al. Immune-mediated regression of established B16F10 melanoma by intratumoral injection of attenuated Toxoplasma gondii protects against rechallenge. Journal of immunology. 2013;190(1):469–78. doi: 10.4049/jimmunol.1201209. PubMed PMID: 23225891; PubMed Central PMCID: PMC3529845.

34. Baird JR, Fox BA, Sanders KL, Lizotte PH, Cubillos-Ruiz JR, Scarlett UK, et al. Avirulent Toxoplasma gondii generates therapeutic antitumor immunity by reversing immunosuppression in the ovarian cancer microenvironment. Cancer research. 2013;73(13):3842–51. doi: 10.1158/0008-5472.CAN-12-1974. PubMed PMID: 23704211; PubMed Central PMCID: PMC3702636.

35. Xu LQ, Yao LJ, Jiang D, Zhou LJ, Chen M, Liao WZ, et al. A uracil auxotroph Toxoplasma gondii exerting immunomodulation to inhibit breast cancer growth and metastasis. Parasites & vectors. 2021;14(1):601. doi: 10.1186/s13071-021-05032-6. PubMed PMID: 34895326; PubMed Central PMCID: PMC8665513.

36. Zhu YC, Elsheikha HM, Wang JH, Fang S, He JJ, Zhu XQ, et al. Synergy between Toxoplasma gondii type I DeltaGRA17 immunotherapy and PD-L1 checkpoint inhibition triggers the regression of targeted and distal tumors. Journal for immunotherapy of cancer. 2021;9(11). doi: 10.1136/jitc-2021-002970. PubMed PMID: 34725213; PubMed Central PMCID: PMC8562526.

37. Ching XT, Fong MY, Lau YL. Evaluation of Immunoprotection Conferred by the Subunit Vaccines of GRA2 and GRA5 against Acute Toxoplasmosis in BALB/c Mice. Frontiers in microbiology. 2016;7:609. doi: 10.3389/fmicb.2016.00609. PubMed PMID: 27199938; PubMed Central PMCID: PMC4847622.

38. Fox BA, Bzik DJ. Nonreplicating, cyst-defective type II Toxoplasma gondii vaccine strains stimulate protective immunity against acute and chronic infection. Infection and immunity. 2015;83(5):2148–55. doi: 10.1128/IAI.02756-14. PubMed PMID: 25776745; PubMed Central PMCID: PMC4399033.

39. Zhang H, Xie W, Zhang Y, Dong X, Liu C, Yi J, et al. Oncolytic adenoviruses synergistically enhance anti-PD-L1 and anti-CTLA-4 immunotherapy by modulating the tumour microenvironment in a 4T1 orthotopic mouse model. Cancer Gene Ther. 2022;29(5):456–65. Epub 2021/09/26. doi: 10.1038/s41417-021-00389-3. PubMed PMID: 34561555; PubMed Central PMCID: PMCPMC9113929.

40. Cui C, Wang J, Fagerberg E, Chen PM, Connolly KA, Damo M, et al. Neoantigen-driven B cell and CD4 T follicular helper cell collaboration promotes anti-tumor CD8 T cell responses. Cell. 2021;184(25):6101–18 e13. Epub 2021/12/02. doi: 10.1016/j.cell.2021.11.007. PubMed PMID: 34852236; PubMed Central PMCID: PMCPMC8671355.

41. Griffith MB, Pearce CS, Heaslip AT. Dense granule biogenesis, secretion, and function in Toxoplasma gondii. The Journal of eukaryotic microbiology. 2022;69(6):e12904. Epub 2022/03/19. doi: 10.1111/jeu.12904. PubMed PMID: 35302693; PubMed Central PMCID: PMCPMC9482668.

42. Braun L, Travier L, Kieffer S, Musset K, Garin J, Mercier C, et al. Purification of Toxoplasma dense granule proteins reveals that they are in complexes throughout the secretory pathway. Molecular and biochemical parasitology. 2008;157(1):13–21. Epub 2007/10/26. doi: 10.1016/j.molbiopara.2007.09.002. PubMed PMID: 17959262.

43. Gendrin C, Mercier C, Braun L, Musset K, Dubremetz JF, Cesbron-Delauw MF. Toxoplasma gondii uses unusual sorting mechanisms to deliver transmembrane proteins into the host-cell vacuole. Traffic (Copenhagen, Denmark). 2008;9(10):1665–80. Epub 2008/07/18. doi: 10.1111/j.1600-0854.2008.00793.x. PubMed PMID: 18631244.

44. Nadipuram SM, Thind AC, Rayatpisheh S, Wohlschlegel JA, Bradley PJ. Proximity biotinylation reveals novel secreted dense granule proteins of Toxoplasma gondii bradyzoites. PloS one. 2020;15(5):e0232552. Epub 2020/05/07. doi: 10.1371/journal.pone.0232552. PubMed PMID: 32374791; PubMed Central PMCID: PMCPMC7202600.

45. Buchholz KR, Bowyer PW, Boothroyd JC. Bradyzoite pseudokinase 1 is crucial for efficient oral infectivity of the Toxoplasma gondii tissue cyst. Eukaryotic cell. 2013;12(3):399–410. Epub 2013/01/08. doi: 10.1128/ec.00343-12. PubMed PMID: 23291621; PubMed Central PMCID: PMCPMC3629768.

46. Guiton R, Vasseur V, Charron S, Arias MT, Van Langendonck N, Buzoni-Gatel D, et al. Interleukin 17 receptor signaling is deleterious during Toxoplasma gondii infection in susceptible BL6 mice. The Journal of infectious diseases. 2010;202(3):427–35. Epub 2010/06/26. doi: 10.1086/653738. PubMed PMID: 20575661.

47. Gaddi PJ, Yap GS. Cytokine regulation of immunopathology in toxoplasmosis. Immunol Cell Biol. 2007;85(2):155–9. Epub 2007/01/18. doi: 10.1038/sj.icb.7100038. PubMed PMID: 17228318.

48. Gazzinelli RT, Wysocka M, Hieny S, Scharton-Kersten T, Cheever A, Kuhn R, et al. In the absence of endogenous IL-10, mice acutely infected with Toxoplasma gondii succumb to a lethal immune response dependent on CD4+ T cells and accompanied by overproduction of IL-12, IFN-gamma and TNF-alpha. Journal of immunology. 1996;157(2):798–805. Epub 1996/07/15. PubMed PMID: 8752931.

49. Daryani A, Hosseini AZ, Dalimi A. Immune responses against excreted/secreted antigens of Toxoplasma gondii tachyzoites in the murine model. Veterinary parasitology. 2003;113(2):123–34. doi: 10.1016/s0304-4017(03)00044-x. PubMed PMID: 12695037.

50. Yuan ZG, Zhang XX, He XH, Petersen E, Zhou DH, He Y, et al. Protective immunity induced by Toxoplasma gondii rhoptry protein 16 against toxoplasmosis in mice. Clinical and vaccine immunology : CVI. 2011;18(1):119–24. doi: 10.1128/CVI.00312-10. PubMed PMID: 21106780; PubMed Central PMCID: PMC3019779.

51. Yuan ZG, Zhang XX, Lin RQ, Petersen E, He S, Yu M, et al. Protective effect against toxoplasmosis in mice induced by DNA immunization with gene encoding Toxoplasma gondii ROP18. Vaccine. 2011;29(38):6614–9. doi: 10.1016/j.vaccine.2011.06.110. PubMed PMID: 21762755.

52. Gold DA, Kaplan AD, Lis A, Bett GC, Rosowski EE, Cirelli KM, et al. The Toxoplasma Dense Granule Proteins GRA17 and GRA23 Mediate the Movement of Small Molecules between the Host and the Parasitophorous Vacuole. Cell host & microbe. 2015;17(5):642–52. doi: 10.1016/j.chom.2015.04.003. PubMed PMID: 25974303; PubMed Central PMCID: PMC4435723.

53. Jongert E, Melkebeek V, De Craeye S, Dewit J, Verhelst D, Cox E. An enhanced GRA1-GRA7 cocktail DNA vaccine primes anti-Toxoplasma immune responses in pigs. Vaccine. 2008;26(8):1025–31. doi: 10.1016/j.vaccine.2007.11.058. PubMed PMID: 18221825.

54. Liu MM, Yuan ZG, Peng GH, Zhou DH, He XH, Yan C, et al. Toxoplasma gondii microneme protein 8 (MIC8) is a potential vaccine candidate against toxoplasmosis. Parasitology research. 2010;106(5):1079–84. doi: 10.1007/s00436-010-1742-0. PubMed PMID: 20177910.

55. Li J, Galon EM, Guo H, Liu M, Li Y, Ji S, et al. PLK:Deltagra9 Live Attenuated Strain Induces Protective Immunity Against Acute and Chronic Toxoplasmosis. Frontiers in microbiology. 2021;12:619335. doi: 10.3389/fmicb.2021.619335. PubMed PMID: 33776955; PubMed Central PMCID: PMC7991750.

56. Yang J, Yang C, Qian J, Li F, Zhao J, Fang R. Toxoplasma gondii alpha-amylase deletion mutant is a promising vaccine against acute and chronic toxoplasmosis. Microbial biotechnology. 2020;13(6):2057–69. doi: 10.1111/1751-7915.13668. PubMed PMID: 32959958; PubMed Central PMCID: PMC7533317.

57. Howe DK, Sibley LD. Toxoplasma gondii comprises three clonal lineages: correlation of parasite genotype with human disease. The Journal of infectious diseases. 1995;172(6):1561–6. doi: 10.1093/infdis/172.6.1561. PubMed PMID: 7594717.

58. Funes SC, Rios M, Escobar-Vera J, Kalergis AM. Implications of macrophage polarization in autoimmunity. Immunology. 2018;154(2):186–95. doi: 10.1111/imm.12910. PubMed PMID: 29455468; PubMed Central PMCID: PMC5980179.

59. Yang L, Zhang Y. Tumor-associated macrophages: from basic research to clinical application. Journal of hematology & oncology. 2017;10(1):58. doi: 10.1186/s13045-017-0430-2. PubMed PMID: 28241846; PubMed Central PMCID: PMC5329931.

60. Glasner A, Levi A, Enk J, Isaacson B, Viukov S, Orlanski S, et al. NKp46 Receptor-Mediated Interferon-gamma Production by Natural Killer Cells Increases Fibronectin 1 to Alter Tumor Architecture and Control Metastasis. Immunity. 2018;48(2):396–8. doi: 10.1016/j.immuni.2018.01.010. PubMed PMID: 29466761; PubMed Central PMCID: PMC5823842.

61. Borst J, Ahrends T, Babala N, Melief CJM, Kastenmuller W. CD4(+) T cell help in cancer immunology and immunotherapy. Nature reviews Immunology. 2018;18(10):635–47. doi: 10.1038/s41577-018-0044-0. PubMed PMID: 30057419.

62. Raskov H, Orhan A, Christensen JP, Gogenur I. Cytotoxic CD8(+) T cells in cancer and cancer immunotherapy. British journal of cancer. 2021;124(2):359–67. doi: 10.1038/s41416-020-01048-4. PubMed PMID: 32929195; PubMed Central PMCID: PMC7853123.

63. Kennedy R, Celis E. Multiple roles for CD4+ T cells in anti-tumor immune responses. Immunological reviews. 2008;222:129–44. doi: 10.1111/j.1600-065X.2008.00616.x. PubMed PMID: 18363998.

64. Kobayashi M, Fitz L, Ryan M, Hewick RM, Clark SC, Chan S, et al. Identification and purification of natural killer cell stimulatory factor (NKSF), a cytokine with multiple biologic effects on human lymphocytes. The Journal of experimental medicine. 1989;170(3):827–45. doi: 10.1084/jem.170.3.827. PubMed PMID: 2504877; PubMed Central PMCID: PMC2189443.

65. Stern AS, Podlaski FJ, Hulmes JD, Pan YC, Quinn PM, Wolitzky AG, et al. Purification to homogeneity and partial characterization of cytotoxic lymphocyte maturation factor from human B-lymphoblastoid cells. Proceedings of the National Academy of Sciences of the United States of America. 1990;87(17):6808–12. doi: 10.1073/pnas.87.17.6808. PubMed PMID: 2204066; PubMed Central PMCID: PMC54627.

66. Wang JL, Elsheikha HM, Zhu WN, Chen K, Li TT, Yue DM, et al. Immunization with Toxoplasma gondii GRA17 Deletion Mutant Induces Partial Protection and Survival in Challenged Mice. Frontiers in immunology. 2017;8:730. Epub 2017/07/15. doi: 10.3389/fimmu.2017.00730. PubMed PMID: 28706518; PubMed Central PMCID: PMCPMC5489627.

67. Gazzinelli RT, Wysocka M, Hayashi S, Denkers EY, Hieny S, Caspar P, et al. Parasite-induced IL-12 stimulates early IFN-gamma synthesis and resistance during acute infection with Toxoplasma gondii. Journal of immunology. 1994;153(6):2533–43. PubMed PMID: 7915739.

68. Suzuki Y, Conley FK, Remington JS. Importance of endogenous IFN-gamma for prevention of toxoplasmic encephalitis in mice. Journal of immunology. 1989;143(6):2045–50. PubMed PMID: 2506275.

69. Sasai M, Yamamoto M. Innate, adaptive, and cell-autonomous immunity against Toxoplasma gondii infection. Experimental & molecular medicine. 2019;51(12):1–10. doi: 10.1038/s12276-019-0353-9. PubMed PMID: 31827072; PubMed Central PMCID: PMC6906438.

70. Yarovinsky F. Innate immunity to Toxoplasma gondii infection. Nature reviews Immunology. 2014;14(2):109–21. doi: 10.1038/nri3598. PubMed PMID: 24457485.

71. Plattner F, Yarovinsky F, Romero S, Didry D, Carlier MF, Sher A, et al. Toxoplasma profilin is essential for host cell invasion and TLR11-dependent induction of an interleukin-12 response. Cell host & microbe. 2008;3(2):77–87. doi: 10.1016/j.chom.2008.01.001. PubMed PMID: 18312842.

72. Scanga CA, Aliberti J, Jankovic D, Tilloy F, Bennouna S, Denkers EY, et al. Cutting edge: MyD88 is required for resistance to Toxoplasma gondii infection and regulates parasite-induced IL-12 production by dendritic cells. Journal of immunology. 2002;168(12):5997–6001. doi: 10.4049/jimmunol.168.12.5997. PubMed PMID: 12055206.

73. Yarovinsky F, Zhang D, Andersen JF, Bannenberg GL, Serhan CN, Hayden MS, et al. TLR11 activation of dendritic cells by a protozoan profilin-like protein. Science. 2005;308(5728):1626-9. doi: 10.1126/science.1109893. PubMed PMID: 15860593.

74. Fox BA, Sanders KL, Chen S, Bzik DJ. Targeting tumors with nonreplicating Toxoplasma gondii uracil auxotroph vaccines. Trends in parasitology. 2013;29(9):431–7. doi: 10.1016/j.pt.2013.07.001. PubMed PMID: 23928100; PubMed Central PMCID: PMC3777737.

75. Sanders KL, Fox BA, Bzik DJ. Attenuated Toxoplasma gondii therapy of disseminated pancreatic cancer generates long-lasting immunity to pancreatic cancer. Oncoimmunology. 2016;5(4):e1104447. doi: 10.1080/2162402X.2015.1104447. PubMed PMID: 27141388; PubMed Central PMCID: PMC4839330.

76. Dubey JP, Laurin E, Kwowk OC. Validation of the modified agglutination test for the detection of Toxoplasma gondii in free-range chickens by using cat and mouse bioassay. Parasitology. 2016;143(3):314–9. Epub 2015/12/03. doi: 10.1017/S0031182015001316. PubMed PMID: 26625933.

